# Cardiac Reprogramming with Drug Resistance Alleviates Doxorubicin-induced Cardiotoxicity in Mice and Pigs

**DOI:** 10.1101/2024.11.28.625834

**Authors:** Yixin Zhang, Weirun Li, Yingxian Xiao, Li Luo, Jiacong Ai, Shushan Mo, Lanya Li, Junyao Deng, Xueyi Wang, Qishan Li, Yan Zeng, Huifang Liu, Fei Wang, Zhenhua Li

## Abstract

Doxorubicin (Dox)-induced cardiotoxicity represents a significant clinical adverse effect associated with cancer chemotherapy treatment. The systematic toxicity associated with the currently available drug dexrazoxane has limited its clinical utility. Inspired by cancer drug resistance mechanisms, we propose a novel strategy termed **T**ransient **o**verexpression of **p**-glycoprotein (P-gp) for **Ca**rdiac **re**programming (TopCare) to induce cardiac drug resistance as a treatment for cardiotoxicity. This approach involves reprogramming cardiomyocytes by delivering lipid nanoparticles (LNPs)-based mRNA therapeutics to induce temporary P-gp overexpression, which in turn reduces intracellular Dox levels and suppresses cytotoxic effects. Our results demonstrated that LNPs with P-gp mRNAs (P-gp LNPs) enhance P-gp expression in H9c2 cells, significantly lowering the cytotoxicity of both Dox and paclitaxel (PTX). In a mouse model with Dox-induced cardiotoxicity, it was observed that P-gp LNPs effectively promoted P-gp overexpression in cardiomyocytes, with a time-dependent decline in P-gp protein levels. This TopCare strategy resulted in improved survival rates, restored cardiac function, and reduced myocardial fibrosis and structural cardiac alterations. Furthermore, studies in large animals have shown that intrapericardial (iPC) injection of P-gp LNPs effectively mitigates adverse effects and restores cardiac function in pig models of Dox-induced cardiotoxicity. The significant cardioprotective effects achieved through cardiac drug resistance highlight the safety, efficacy, and clinical potential of the TopCare strategy for alleviating doxorubicin-induced cardiotoxicity.

## Introduction

Chemotherapy remains a principal therapeutic modality for cancer, offering significant benefits in symptom management, prolonging survival, and enhancing patients’ quality of life(1). Dox is a frontline chemotherapy drug included in the WHO Model List of Essential Medicines and is widely utilized to treat breast cancer, leukaemia, lymphoma and other cancers(2–4). However, the damage that Dox causes to healthy tissue represents a significant limitation to its clinical application(5). In addition to the typical systemic toxicity, Dox is demonstrated to be associated with serious cardiomyopathy in a dose-dependent manner(6, 7). A significant body of evidence indicates that 8.8% of cancer survivors die from cardiovascular disease, with as many as a quarter of patients exhibiting Dox-induced cardiotoxicity. Dox causes cardiomyocyte death by the generation of a high level of reactive oxygen species (ROS) and accumulating of iron, which subsequently lead to lipid peroxidation-dependent death(8). This results in pathological cardiomyopathy, characterized by cytoplasmic vacuolization of the heart and thinning and disruption of myogenic fibers(9). At present, the only FDA-approved pharmaceutical drug for the treatment of Dox-induced cardiotoxicity is dexrazoxane, which is an iron chelator that inhibits the effects of lipid peroxidation and cardiomyocyte damage caused by Dox(10). However, this agent has been associated with adverse effects, including altered liver function and hematological toxicity(11). It is therefore imperative to develop an alternative safe and efficacious method for alleviating Dox-induced cardiotoxicity.

As the primary target of chemotherapeutic drugs, cancer cells usually develop drug resistance through different mechanisms, with the intrinsic or adaptive overexpression of drug efflux proteins representing a significant contributing factor(12–14). The development of drug resistance contributes to reduced cancer cell-killing efficacy of chemotherapeutic agents(15). A well-studied drug transporter protein is P-gp, also known as MDR1 or ABCB1, which is a key ATP-binding cassette transporter(16, 17). It is a cell membrane glycoprotein that is commonly overexpressed in multidrug-resistant tumor cells and facilitates the transport of intracellular chemotherapeutic drugs across the membrane and out of the cell(18, 19). Prior studies have demonstrated that the anti-cancer effects can be significantly enhanced by the co-delivery of P-gp inhibitors and Dox, which results in the down-regulated expression of P-gp and subsequent enhancement of Dox accumulation(20, 21). Conversely, it could be postulated that the overexpression of P-gp functions to protect the cells from chemotherapeutic drugs(22, 23). By learning from the cancer drug resistance, we propose that the reprogramming of cardiomyocytes and the establishment of “chemotherapeutic drug resistance in the heart” (cardiac drug resistance) by enhancing the cardiac P-gp expression might be an efficient approach to alleviate the Dox-induced cardiotoxicity.

The overexpression of proteins can be achieved through the administration of messenger RNA (mRNA) and DNA, as well as through the stimulation of signaling pathways(3, 24). It is noteworthy that mRNA serves as a transient carrier of genetic information, which is then is translated into protein(25–27). The absence of nuclear entry and genome integration for mRNA reduces the risk of insertion mutagenesis and oncogenic effects, thereby enhancing the safety and clinical applicability of mRNA therapeutics(28). Furthermore, in comparison to DNA or virus-based gene regulation, the processes involved in the production of mRNA are more straightforward, less costly and easier to scale up to industrial levels of production(29–32). In recent decades, mRNA-based therapeutics have been widely employed in the treatment of various diseases, with numerous instances demonstrating encouraging efficacy in preclinical studies and even clinical applications(27, 33–35). Therefore, the overexpression of P-gp in the heart using mRNA-based therapeutics represents an attractive strategy.

In this study, we developed a TopCare strategy by learning from cancer drug resistance and establishing cardiac drug resistance. In brief, P-gp mRNA was administered to reprogram cardiomyocytes, leading to P-gp overexpression in the heart and providing cardiac protection against chemotherapeutic agents. The in vitro experiments demonstrated that the administration of P-gp LNPs was capable of successfully overexpressing P-gp in cardiomyocytes, reducing the intracellular Dox content and decreasing Dox cytotoxicity. In vivo analysis revealed that intramyocardial administration of P-gp LNPs resulted in transient overexpression of P-gp in the cardiac tissue, with the protein returning to normal levels within four days. The results of mouse studies demonstrated that P-gp LNPs could effectively overexpress P-gp in the mouse heart, improve the survival rate, and enhance the cardiac function in the mice of Dox-induced cardiotoxicity, and mitigate the myocardial fibrosis and cardiac structural disorders induced by Dox. Furthermore, large animal experiments demonstrated that P-gp LNPs effectively attenuated the adverse effects of Dox on cardiac function. This study reports a safe and effective TopCare approach for alleviating chemotherapy-induced cardiotoxicity through the establishment of cardiac drug resistance via transient overexpression of P-gp.

## Results

### In vitro P-gp reprogramming of cardiomyocytes

Lipid nanoparticles (LNPs) have been considered an effective and secure approach for the administration of mRNA-based therapeutic drugs, with LNP-mRNA entering clinical trials and applications for the treatment of various diseases(36, 37). In this study, we prepared four-component LNPs for the encapsulation and delivery of P-gp mRNAs. The intrapericardial (iPC) delivery of P-gp mRNAs via LNPs are expected to result in the transient overexpression of P-gp in the heart, thereby accelerating the efflux of doxorubicin (Dox) out of cardiomyocytes and protecting the heart from Dox-induced cardiotoxicity (Fig. 1a). First, the morphology and size distribution of LNPs encapsulating P-gp mRNAs (P-gp LNPs) were determined using transmission electron microscopy (TEM) imaging and dynamic light scattering (DLS) analysis. As demonstrated, the P-gp LNPs were observed to be monodisperse and spherical (Supplementary Fig. S1a). The average radius was measured to be 115 nm with a polydispersity index (PDI) of 0.174 (Supplementary Fig. S1b). The encapsulation efficiency of mRNAs in LNPs was shown to be 92.21%. CCK-8 analysis with H9c2 cells showed that the prepared LNPs and P-gp LNPs exhibited no cytotoxic effects at a high concentration (Supplementary Fig. S2a). Furthermore, no obvious hemolytic effect was observed when mouse blood samples were incubated with P-gp LNPs treatment even at a high concentration (Supplementary Fig. S2b,c), indicating the high biocompatibility of the LNPs and P-gp LNPs. We then incubated H9c2 cells with P-gp LNPs to evaluate their in vitro transfection efficiency. Western blot results demonstrated a significant up-regulation in P-gp protein expression in H9c2 cells treated with P-gp LNPs compared to the control group (Fig. 1b,c). Increased P-gp expression was also observed in the immunofluorescence staining and flow cytometry analysis (Fig. 1d and Supplementary Fig. S3a). In addition, the western blotting analysis and flow cytometry analyses revealed that the overexpression of P-gp protein in the cardiomyocytes reached its peak on the first day following the treatment of P-gp LNPs, and gradually declined over time (Fig. 1e,f and Supplementary Fig. S3b). The expression of P-gp was demonstrated to reduce to its initial normal level on the 7th day after P-gp LNP incubation. These results demonstrated the in vitro effective, transient reprogramming of cardiomyocytes with P-gp overexpression by using P-gp LNPs. Given the nature of interval treatment of chemotherapy, the transient reprogramming of cardiomyocytes via mRNA is an attractive approach for Dox-induced cardiotoxicity due to the precise dosing and limited side effects(38).

**Fig. 1.**
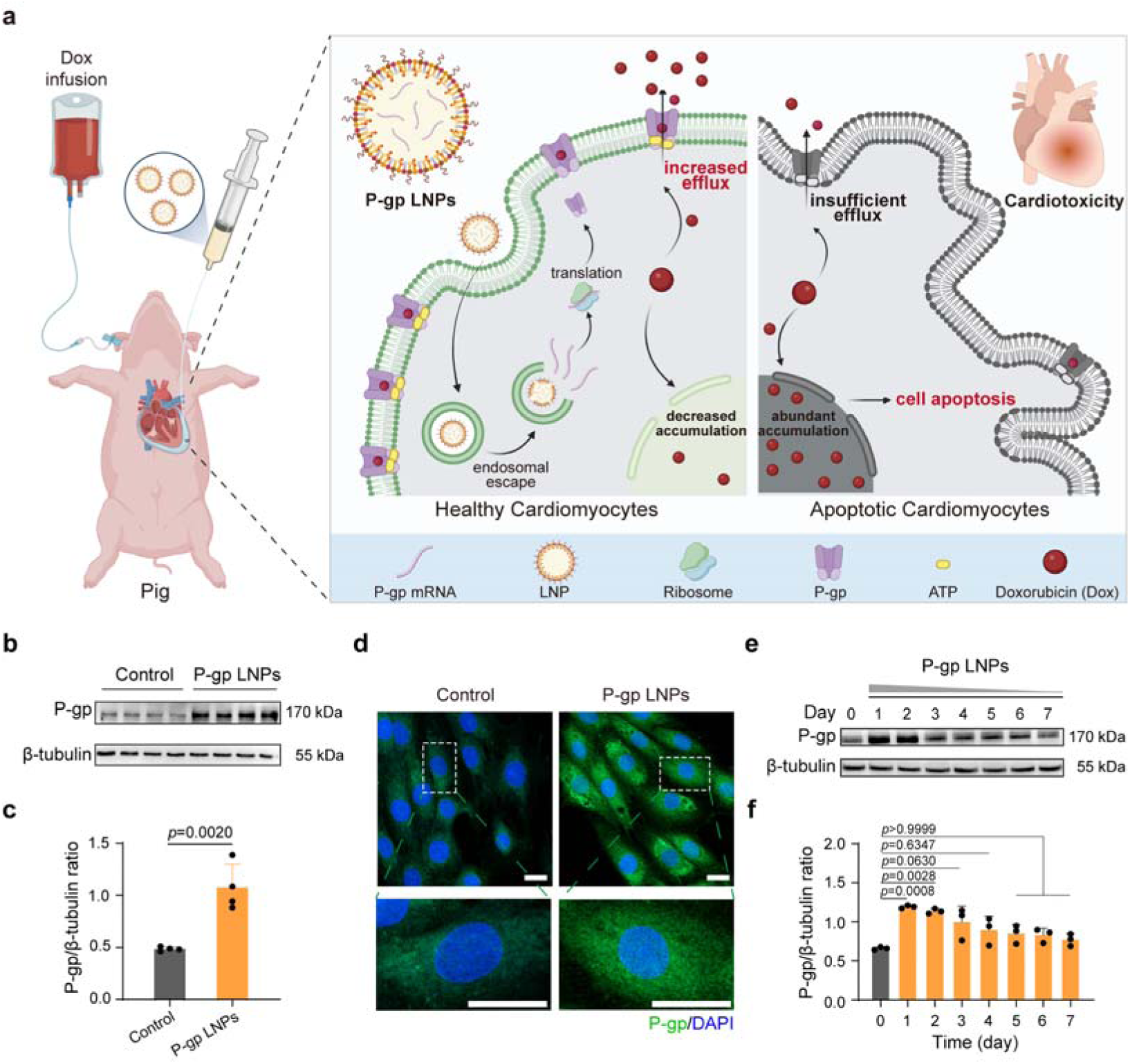
In vitro transient P-gp reprogramming of H9c2 cells using P-gp LNPs. **a**, Schematic illustration of the establishment of cardiac drug resistance in pigs via P-gp reprogramming to alleviate Dox-induced cardiotoxicity. P-gp LNPs were administered via intrapericardial (iPC) injection, followed by Dox administered via the ear vein 24 h later. P-gp LNPs lead to the overexpression of P-gp in the cardiomyocytes, thereby reducing the accumulation of Dox in the cell and alleviating the cardiotoxicity. **b,c**, Western blotting (**b**) and statistical analysis (**c**) of P-gp expression in H9c2 cells with indicated treatments (Control, P-gp LNPs) (*n* = 4 independent biological samples). **d**, Representative immunofluorescence staining images of P-gp (green) overexpression in H9c2 cells incubated with P-gp LNPs for 24 h, with a magnified inset. DAPI staining was employed to visualize the nucleus (blue). Scale bars, 20 μm (top); 20 μm (bottom). Experiments were replicated in triplicate. **e,f**, Western blotting analysis (**e**) and statistical analysis (**f**) of the attenuation of the overexpressed P-gp in H9c2 cells after incubation with P-gp LNPs over a 7-day period (*n* = 3 independent biological samples). Significant differences were determined by two-tailed unpaired Student’s *t*-test (**c**), or one-way ANOVA and Bonferroni’s multiple comparisons test (**f**). Results are shown as meanL±Ls.d. Schematic in **a** and pig cartoon were created with BioRender.com.

### P-gp reprogramming facilitates Dox efflux in cardiomyocytes

As one of the identified primary inducers of cancer drug resistance, P-glycoprotein (P-gp) is able to facilitate the transport of intracellular chemotherapeutic drugs, such as Dox, across the cell membrane and out of the cell(39). We therefore investigated the effect of P-gp reprogramming induced by P-gp LNPs on the intracellular Dox levels in cardiomyocytes (Fig. 2a). As shown, the confocal laser scanning microscopy (CLSM) imaging and corresponding fluorescence intensity analysis revealed strong red fluorescence (approximately 150 a.u.) in the nucleus of H9c2 cells after 24 h or 36 h incubation with Dox, indicating a high level of Dox in the cells. In contrast, a relatively lower fluorescence signal of Dox (approximately 100 and 50 a.u.) was observed in H9c2 cells that pre-treated with P-gp LNPs (Fig. 2b-d). Notably, the H9c2 cells treated with Dox maintained a high level of intracellular Dox at 36 h compared to 24 h, whereas the H9c2 cells pre-treated with P-gp LNPs showed a marked reduction. (Fig. 2c,d). Flow cytometry analysis also confirmed the significant reduction of Dox levels in the H9c2 cells following P-gp LNPs treatment, with a mean fluorescence intensity decreasing from approximately 2729 to 1061 (Fig. 2e,f). These results demonstrated that P-gp LNPs treatment could significantly reduce the intracellular concentration of Dox, which can be attributed the rapid and continuous efflux of Dox from cells induced by P-gp overexpression in the cardiomyocytes. In addition, we investigated the impact of P-gp LNPs treatment on the Dox-associated cytotoxicity. In vitro studies revealed that pre-treatment with P-gp LNPs markedly increased the cell viability of H9c2 cells with Dox treatment at various concentrations (Fig. 2g), indicating that P-gp overexpression protects the cardiomyocytes against the Dox-induced cytotoxicity. Since P-gp has been demonstrated to facilitate the transport of a range of chemotherapeutic agents, including paclitaxel (PTX)(40), which could also induce cardiotoxicity when employed in chemotherapy(41, 42), we also examined the impact of P-gp reprogramming on paclitaxel (PTX) in H9C2 cells. As shown, P-gp overexpression via P-gp LNPs impeded the mitotic arrest caused by PTX and markedly attenuated the PTX-induced cytotoxicity in the cardiomyocytes (Supplementary Fig. S4).

**Fig. 2.**
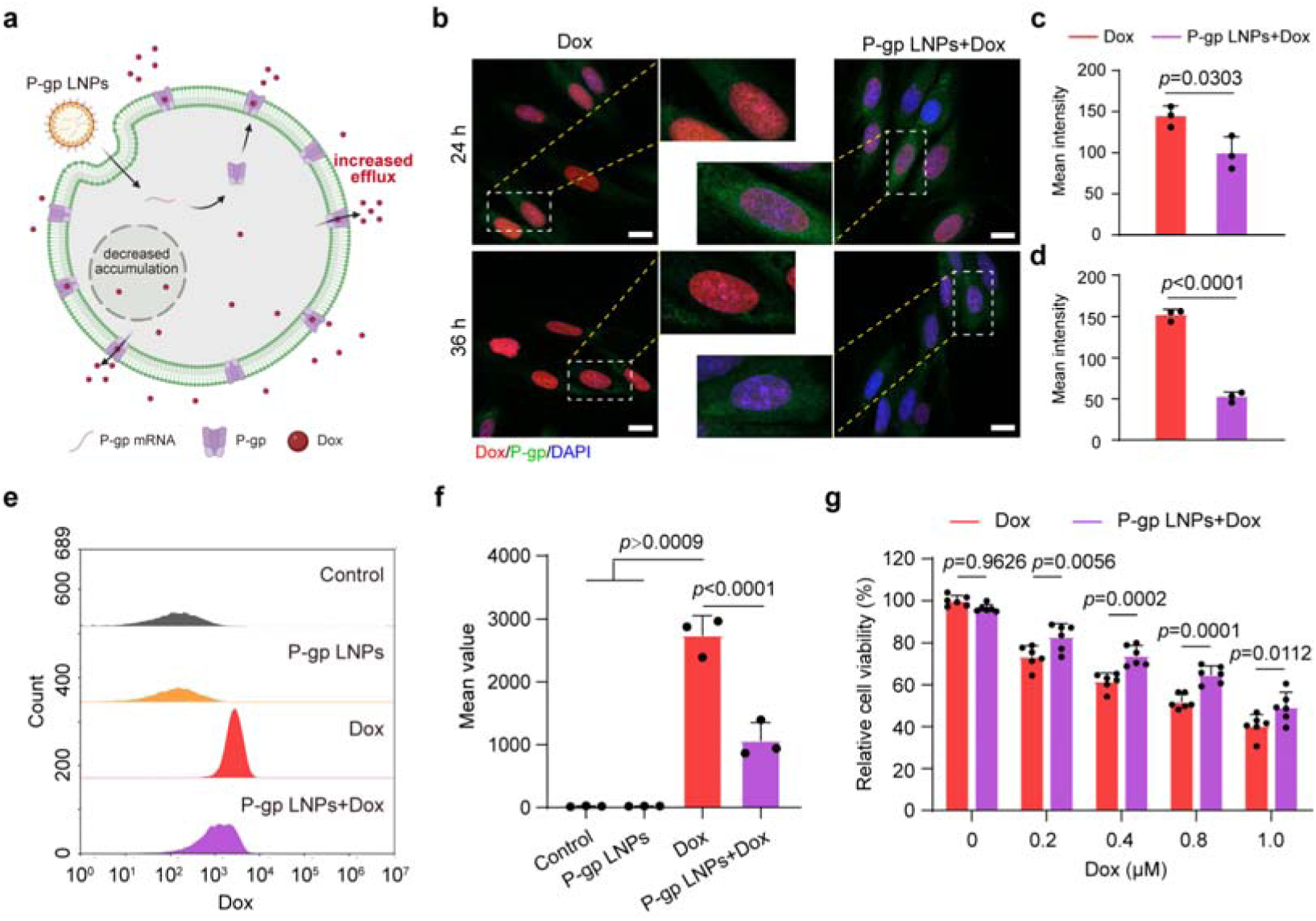
In vitro P-gp overexpression reduced intracellular Dox in cardiomyocytes. **a,** Schematic illustration of the P-gp reprogramming via P-gp LNPs to increase the Dox efflux in cardiomyocytes. **b**, Representative immunofluorescence staining images of H9c2 cells with the indicated treatments (Dox and P-gp LNPs+Dox). Cells were treated with a 24 h pretreatment of PBS or P-gp LNPs prior to incubation with Dox for 24 h or 36 h. P-gp was stained green. DAPI staining was employed to visualize the nucleus (blue). Scale bar, 20 μm. Experiments were replicated in triplicate. **c,d**, Quantified data of the Dox signal in H9c2 cells incubated with Dox for 24 h (**c**) or 36 h (**d**). **e,f**, Flow cytometry analysis (**e**) and quantitative mean value (**f**) of intracellular Dox in H9c2 cells with indicated treatments (Control, P-gp LNPs, Dox, and P-gp LNPs+Dox) (*n* = 3 independent biological samples). **g**, Relative cell viability of H9c2 cells with indicated treatments (Dox and P-gp LNPs+Dox) determined via CCK-8 assays (*n* = 6 independent biological samples). Significant differences were determined by two-tailed unpaired Student’s *t*-test (**c,d**), one-way ANOVA and Bonferroni’s multiple comparisons test (**f**), or two-way ANOVA and Bonferroni’s multiple comparisons test (**g**). Results are shown as meanL±Ls.d.

### In vivo LNP-mRNA biodistribution and cardiac P-gp reprogramming

We next investigated the biodistribution of the P-gp LNPs after intramyocardial delivery and the in vivo cardiac P-gp reprogramming efficacy in mice. To determine the biodistribution of LNP-mRNA, LNPs encapsulating luciferase mRNAs (Luc LNPs) were intramyocardially injected into the myocardium of B16F10 tumor-bearing mice for imaging (Fig. 3a). As shown, the results of the In Vivo Imaging Systems (IVIS) demonstrated strong luminescence intensity in the cardiac region, whereas no detectable luminescence was observed at the B16F0 tumor site (Fig. 3b). Ex vivo imaging of harvested organs also showed a dominant distribution of luminescence in the heart and no signal in the tumor tissue (Fig. 3c,d). In addition, the immunostaining results indicated the overexpression of the luciferase in the heart, but no alteration in the luciferase expression level was observed in the tumor (Supplementary Fig. S5). We further studied the precise localization of the proteins in the cardiac tissue after mRNA translation using LNPs encapsulating eGFP mRNAs (eGFP LNPs). The immunostaining results showed the dominant distribution of eGFP proteins in the left ventricular myocardial tissue as evidenced by the colocalization of eGFP with α-SA (Fig. 3e). In contrast, the rare colocalization of α-SMA and eGFP in the left ventricle suggested that only a limited portion of the blood vessels were transfected by the eGFP LNPs (Fig. 3f). These results collectively proved the ideal biodistribution of LNP-mRNA in the cardiomyocytes after the intramyocardial delivery, as the Dox-induced cardiotoxicity is characterized by the initial myocardial cell damage and subsequent left ventricular dysfunction(43, 44). The in vivo biocompatibility of P-gp LNPs was assessed prior to in vivo application of P-gp LNPs in cardiac reprogramming. The initial administration of P-gp LNPs was conducted intramyocardially in healthy mice. After a seven-day observation period, the mice were sacrificed to harvest of major organs and blood samples. As shown in the haematoxylin and eosin (H&E) images, no notable morphological abnormalities were detected in main organs of the mice treated with P-gp LNPs (Supplementary Fig. S6).

**Fig. 3.**
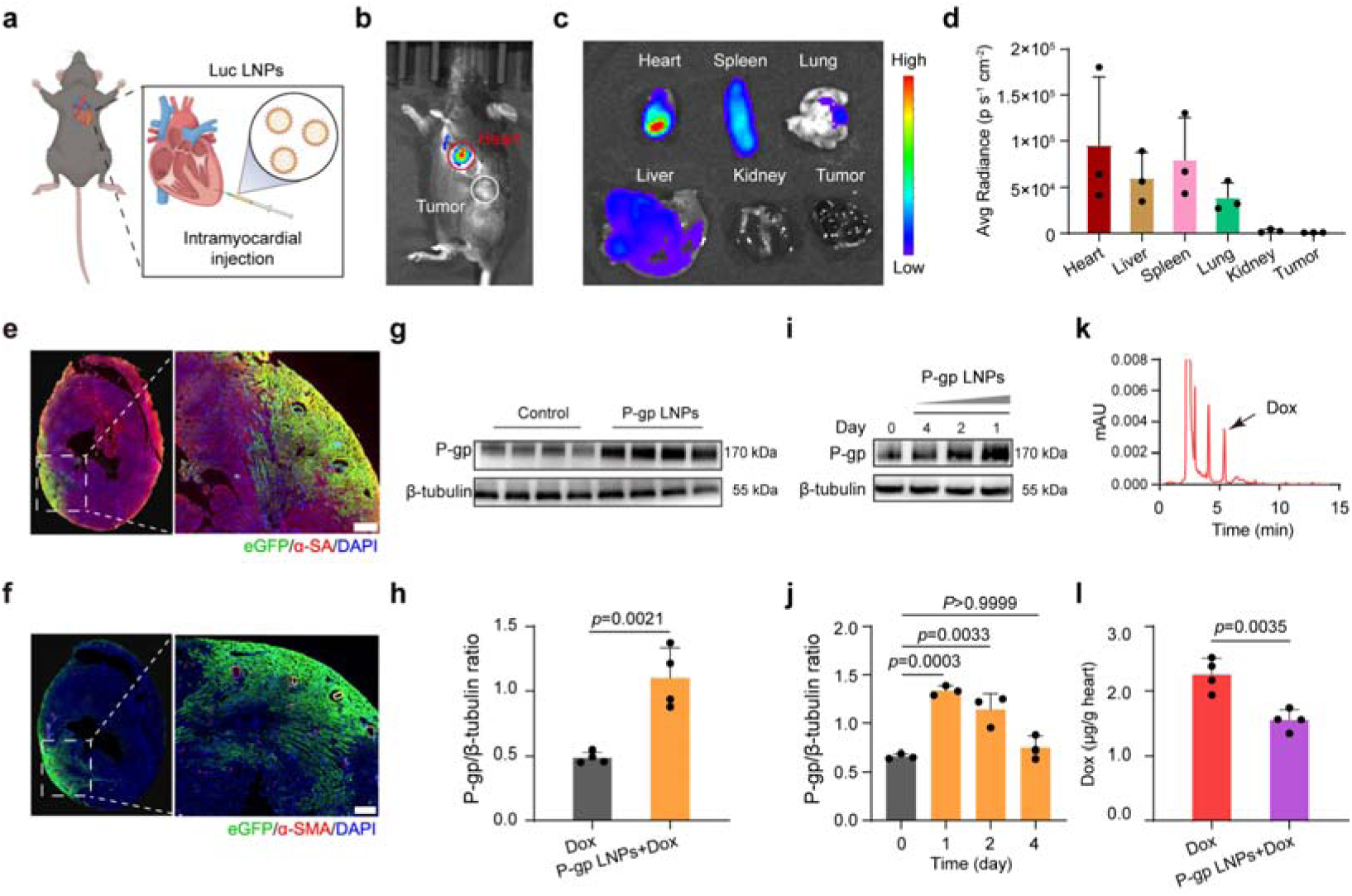
In vivo biodistribution of Luc LNPs and cardiac P-gp reprogramming with drug resistance. **a**, Schematic illustration of the intramyocardial injection of Luc LNPs in C57BL/6 mice. **b**, Representative in vivo bioluminescence imaging of B16F10 tumor-bearing mouse 24 h after intramyocardial injection of Luc LNPs. Experiments were replicated in triplicate. **c,d**, Representative ex vivo bioluminescence imaging (**c**) and quantitative analysis (**d**) of harvested organs (heart, spleen, lung, liver, kidneys, and tumor) in B16F10 tumor-bearing mice (*n* = 3 biologically independent mice in each group). **e,f**, Representative immunofluorescence staining of GFP^+^ (green) myocytes at the injection site 24 h post injection of eGFP LNPs (left) with magnified inset (right). α-SA (**e**) or α-SMA (**f**) was stained red. DAPI staining was employed to visualize the nucleus (blue). Scale bar, 200 μm. Experiments were replicated in triplicate. **g,h**, Western blotting (**g**) and statistical analysis (**h**) of the P-gp expression in the hearts of mice with indicated treatments (Control, P-gp LNPs) (*n* = 4 independent biological mice in each group). **i,j**, Western blotting (**i**) and statistical analysis (**j**) of the attenuation of the overexpressed P-gp in the hearts of mice after incubation with P-gp LNPs over a 4-day period (*n* = 3 independent biological mice per group). **k,l**, High-performance liquid chromatography (HPLC) analysis (**k**) and quantitative analysis (**l**) of Dox in the mice hearts (*n* = 4 independent biological mice per group). Significant differences were determined by two-tailed unpaired Student’s *t*-test (**h,l**) or one-way ANOVA and Bonferroni’s multiple comparisons test (**j**). Results are shown as meanL±Ls.d. Schematic in **a** and mouse cartoon were created with BioRender.com.

In addition, no significant differences were detected in the hematological parameters and biochemical indexes, particularly those pertaining to cardiac, hepatic, and renal functions (Supplementary Fig. S7). Of note, the ELISA analysis revealed no significant elevation in proinflammatory TNF-α and IL-6 cytokines following the administration of the P-gp LNPs (Supplementary Fig. S8). We then carried out the cardiac P-gp reprogramming using the established strategy of intramyocardial delivery of therapeutic LNP-mRNA. Western blotting results revealed the cardiac overexpression of P-gp proteins 24 h after a single intramyocardial injection of P-gp LNPs (10 μg mRNA/20 g) (Fig. 3g,h). Instead, the intramyocardial administration of P-gp LNPs resulted in insignificant changes in the tumor P-gp protein levels (Supplementary Fig. S9). Moreover, a time-dependent decrease in P-gp expression was detected and the P-gp levels returned to normal by day 4 after P-gp LNPs administration (Fig. 3i,j). Consequently, high-performance liquid chromatography (HPLC) demonstrated that the pre-treatment of P-gp LNPs resulted in a remarkable decrease of Dox levels in the heart, demonstrating the development of cardiac drug resistance (Figure 3k,l and Supplementary Fig. S10).

### P-gp LNPs alleviate Dox-induced cardiotoxicity in mice

Next, we verified the cardioprotective effect of cardiac P-gp reprogramming via P-gp LNPs against Dox-induced cardiotoxicity in a Dox-challenged mouse model. In brief, P-gp LNPs (10 μg mRNA/20g) were administrated into the cardiac tissue of mice via intramyocardial injection, and the Dox (15 mg/kg) was administrated intraperitoneally, followed by subsequent therapeutic evaluation (Fig. 4a). As shown, the Dox administration resulted in a notable reduction in the mice’s body weight and a 50% decrease in survival rate of the mice over a one month period, indicating the serve systematic toxicity of Dox to the mice (Supplementary Fig. S11a,b). Encouragingly, pre-treatment with P-gp LNPs significantly reversed the body weight loss observed in mice that received Dox treatment. In addition, the survival rate of mice pre-treated with P-gp LNPs was shown to be markedly higher (80%) than that observed in mice treated with Dox. In particular, we calculated heart weight/tibia length (HW/TL) values to evaluate changes in cardiac structure, and found that the P-gp LNPs pre-treatment significantly increased the HW/TL value compared to the Dox-treated group (Supplementary Fig. S11c). Subsequently, we performed echocardiographic analysis to determine the cardiac structure and function at the end of the therapeutic period (Fig. 4b). The ejection fraction (EF) and fractional shortening (FS) of mice that received Dox treatment were severely reduced, indicating that mice received Dox treatment suffered from severely damaged cardiac function (Fig. 4c,d). In contrast, mice in the P-gp LNPs-treated group showed significantly improved EF and FS. Furthermore, a rescued reduction in left ventricular end diastolic diameter (LVIDd) and left ventricular end systolic diameter (LVIDs) was observed in mice pre-treated with P-gp LNPs compared to Dox-treated mice, showing the role of P-gp LNPs in reversing the Dox-induced cardiac remodeling (Fig. 4e,f). Histopathological staining of cardiac tissue sections was used to provide further insight into the cardiac structure and cardio-therapeutic effects of the cardiac reprogramming by P-gp LNPs. We conducted Masson’s trichrome staining to determine the damaged fibrotic area in the heart tissue. As demonstrated, Dox administration induced increased fibrosis in the cardiac tissue of the mice. The mice in the Dox+P-gp LNPs group demonstrated a reduction in myocardial fibrosis in the heart tissue (Fig. 4g,h). Wheat germ agglutinin (WGA) analysis revealed a significant reduction in the cardiac cross-sectional area after Dox administration, indicating the severe cardiomyocyte atrophy in the myocardium. The pre-treatment of P-gp LNPs was shown to remarkably alleviate the cardiomyocyte atrophy induced by Dox treatment (Fig. 4i,j). Furthermore, the TdT-mediated dUTP-biotin nick end labeling (TUNEL) assays showed a notable elevation in apoptotic cells in the myocardium of Dox-challenged mice, which was effectively mitigated by the P-gp LNPs (Supplementary Fig. S12a). Similarly, H&E staining results showed a distinct pathological abnormality in the myocardium of mice treated with Dox, demonstrating the Dox-induced tissue damage in the cardiac tissue sections (Supplementary Fig. S12b). Conversely, the administration of P-gp LNPs was observed to maintain the normal morphological structures of the cardiac tissue. Together, our data suggested that the cardiac P-gp reprogramming with drug resistance could effectively alleviate Dox-induced cardiotoxicity and protect the cardiac structure and function in mice.

**Fig. 4.**
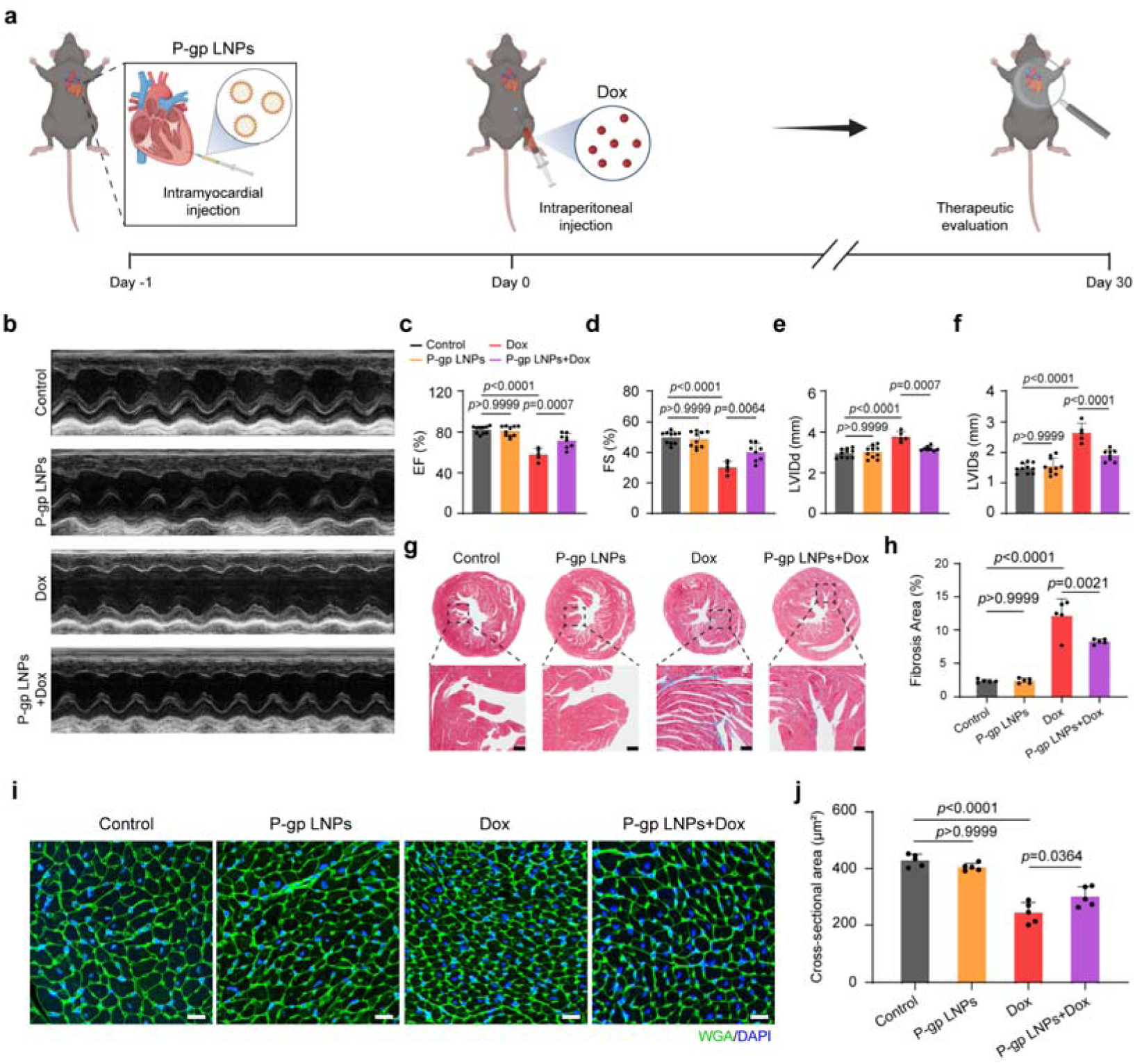
Cardiac P-gp reprogramming improved cardiac function against Dox-induced cardiotoxicity in mice. **a**, Schematic illustration of cardiac P-gp reprogramming by intramyocardial injection of P-gp LNPs and therapeutic evaluation of the cardiac function of Dox-challenged mice. **b-f**, Representative M-mode echocardiographic images (**b**) and statistical analysis of EF, FS, LVIDd, and LVIDs (**c**-**f**) of the mice with indicated treatments (Control, P-gp LNPs, Dox, and P-gp LNPs+Dox) (*n* = 10, 10, 5, and 8 independent biological mice per group, respectively). The therapeutic evaluation of the cardiac function was conducted 30 d after the mice received the indicated treatments. **g,h**, Representative Masson’s trichrome stained images (**g**) and quantitative analysis of the fibrosis area (**h**) of the cardiac tissues in mice with indicated treatments (Control, P-gp LNPs, Dox, and P-gp LNPs+Dox) (n = 5 biologically independent mice in each group). Scale bar, 100 μm. **i,j**, Representative WGA (green) staining images (**i**) and quantitative analysis of cross-sectional area (**j**) in the cardiac tissues from mice with indicated treatments (Control, P-gp LNPs, Dox, and P-gp LNPs+Dox) (n = 5 biologically independent mice per group). DAPI staining was employed to visualize the nucleus (blue). Scale bar, 20 μm. Significant differences were determined by one-way ANOVA and Bonferroni’s multiple comparisons test (**c-f,h,j**). Results are shown as meanL±Ls.d. Schematic in **a** and mouse cartoon were created with BioRender.com.

### Cardiac drug resistance does not compromise in vivo anti-cancer efficacy

Dox is primarily utilized as an anti-cancer chemotherapeutic agent. Thus, the establishment of cardiac drug resistance via P-gp reprogramming in the heart should not compromise the anti-cancer efficacy of Dox therapy. We then conducted in vivo anti-cancer therapy using a B16F10 tumor-bearing mice model. At an average tumor volume of approximately 50 mm^3^, the mice were pre-treated with an intramyocardial injection of P-gp LNPs (10 µg mRNA/20g), and the Dox (15 mg/kg) was administrated intraperitoneally 24 h later, followed by subsequent tumor growth monitoring (Fig. 5a). The results demonstrated that the intramyocardial injection of P-gp LNPs did not suppress tumor growth. Furthermore, both the Dox and Dox+P-gp LNPs treatments demonstrated a notable inhibition of tumor growth in comparison to the control group (Fig. 5b). In addition, the Dox+P-gp LNPs group exhibited a statistically similar degree of tumor growth suppression compared to the Dox group (Fig. 5c, d). Notably, the mice that received Dox treatment exhibited strong systemic side effects, as evidenced by a notable loss of body weight and an elevated mortality rate (Fig. 5e and Supplementary Fig. S13). The systemic toxicity of Dox was mitigated by pre-treating the mice with P-gp LNPs. Histopathologic H&E staining, TUNEL immunofluorescence staining, and Ki67 immunohistochemistry staining demonstrated extensive cancer cell apoptosis and tumor tissue damage in the mice treated with Dox and P-gp LNPs+Dox (Fig. 5f,g and Supplementary Fig. S14). Subsequently, the impact of Dox on cardiac morphology and function during anti-cancer Dox therapy was assessed. As shown, Dox administration resulted in a reduction in HW/TL, EF, and FS, accompanied by an increase in LVIDd and LVIDs. The Dox-induced cardiotoxicity in tumor-bearing mice was also significantly alleviated by the pre-administration of P-gp LNPs (Fig. 5h and Supplementary Fig. S15). Collectively, our results indicated that the establishment of cardiac drug resistance via P-gp LNPs does not induce any inhibitory effects on the anti-tumor effects of Dox when alleviating Dox-induced cardiotoxicity.

**Fig. 5.**
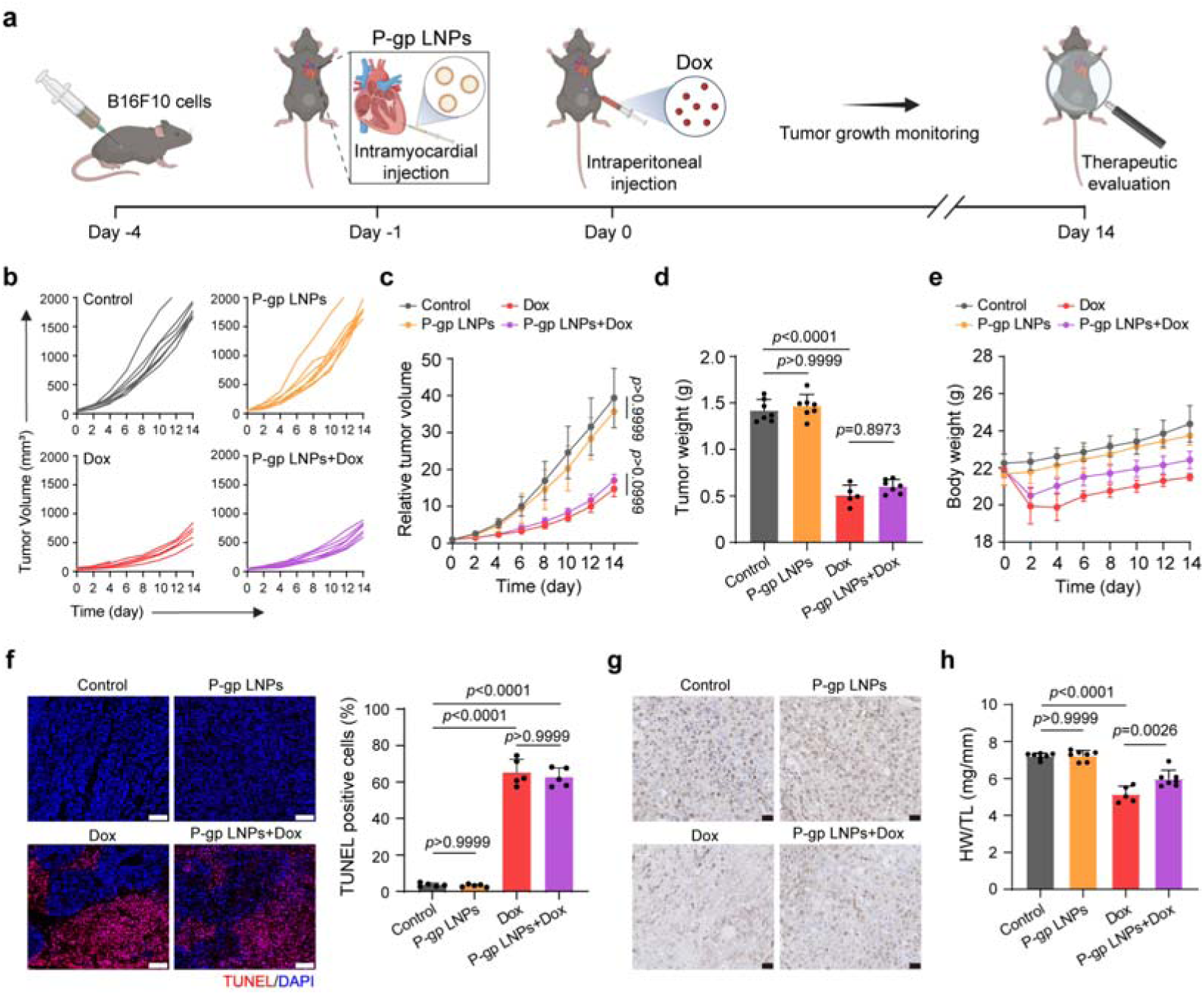
In vivo anti-cancer efficacy and cardioprotective effects in tumor-bearing mice model. **a**, Schematic of the development of B16F10 tumor-bearing mice model for assessment of anti-cancer efficiency and cardioprotective effects after administration of different treatments. **b**, Individual tumor volume changes of mouse in each group with indicated treatments (Control, P-gp LNPs, Dox, and P-gp LNPs+Dox) (*n* = 8 independent biological mice per group). **c**, Relative tumor growth over time in each group with indicated treatments. The tumor growth rate was calculated as the fold change over the initial tumor volume (*V*_t_/*V*_0_) (*n* = 8 independent biological mice per group). **d**, Tumor weight of the mouse in each group with indicated treatments (*n* = 7, 7, 5, and 7 independent biological mice per group, respectively). **e**, Relative body weight changes over time in each group with indicated treatments. The body weight rate was calculated as the fold change over the initial body weight (*W*_t_/*W*_0_) (*n* = 8 independent biological mice per group). **f**, Representative TUNEL immunofluorescence staining images and quantitative analysis of TUNEL positive cells (%) of tumor tissue sections dissected from mice with indicated treatments (Control, P-gp LNPs, Dox, and P-gp LNPs+Dox). (*n* = 5 independent biological mice per group). Scale bar, 100 μm. **g**, Representative Ki67 immunohistochemistry staining images of tumor tissue sections dissected from mice with indicated treatments (Control, P-gp LNPs, Dox, and P-gp LNPs+Dox). (*n* = 5 independent biological mice per group). Scale bar, 20 μm. **h**, HW/TL of the mouse in each group with different treatments (*n* = 7, 7, 5, and 7 independent biological mice per group, respectively). Significant differences were determined by one-way ANOVA and Bonferroni’s multiple comparisons test (**c,d,f,h**). Results are shown as meanL±Ls.d. Schematic in **a** and mouse cartoon were created with BioRender.com.

### Cardiac P-gp reprogramming with drug resistance alleviates Dox-induced cardiotoxicity in Pig

To further investigate the clinical translation potential of our TopCare strategy, we evaluated the therapeutic efficacy of P-gp LNPs in pig studies due to the high similarity between the human and porcine heart organ in terms of size, overall morphology, and immune or physiological activities(45) (Fig. 6a). Similar to the mouse studies, western blotting analysis demonstrated overexpression of P-gp proteins in porcine endothelial (PED) cells after treatment with LNPs encapsulating porcine P-gp mRNA (P-gp LNPs) (Supplementary Fig. S16). The results of flow cytometry experiments revealed a notable reduction in cell fluorescence intensity following P-gp LNPs pretreatment compared to the Dox group, suggesting the successful establishment of drug resistance in the PED cells (Supplementary Fig. S17a,b). In addition, CCK8 assays showed that the P-gp LNPs pretreatment could significantly reduce the cytotoxicity of Dox to PED cells (Supplementary Fig. S17c). Inspired by the in vitro results, we further practiced in a Dox-challenged Bama minipig model for the establishment of cardiac drug resistance. The P-gp LNPs were administrated four times into pig heart via our previously developed mini-invasive iPC injection(46, 47) followed by the Dox (1 mg/kg) infusion (Fig. 6b). The iPC injection is supported by two dependent tiny incisions, one for catheter insertion and the other for endoscope insertion. The catheter was introduced into the pericardial cavity of the porcine heart, followed by the injection of P-gp LNPs (150 µg mRNA/25 kg) under thoracoscopic control (Fig. 6c and Supplementary Movie 1). The echocardiographic results showed that pretreatment with P-gp LNPs can alleviate the Dox-induced decrease in EF and FS, thereby improving the cardiac function of pigs (Fig. 6d). A notable decline in EF and FS was detected in the Dox-challenged pig at both 4 and 12 weeks, suggesting the acute and long-term cardiotoxicity induced by Dox infusion. Pretreatment with P-gp LNPs resulted in a mitigated reduction in EF and FS compared to the Dox-treated pig (Fig. 6e). In addition, multiple staining results of heart tissue sections showed that P-gp LNPs could remarkedly alleviate Dox-induced cardiac fibrosis, cardiomyocyte atrophy, cell apoptosis and pathological abnormality in pig heart (Fig. 6f-j). It can be observed that the T-wave inversion was significantly deeper in the model pigs after 12 weeks of administration, probably due to myocardial injury induced by Dox. The milder T-wave inversion in the P-gp LNPs pretreated group indicated that the cardiac P-gp reprogramming contributed to protect myocardial cells from Dox-induced cardiotoxicity (Supplementary Fig. S18).

**Fig. 6.**
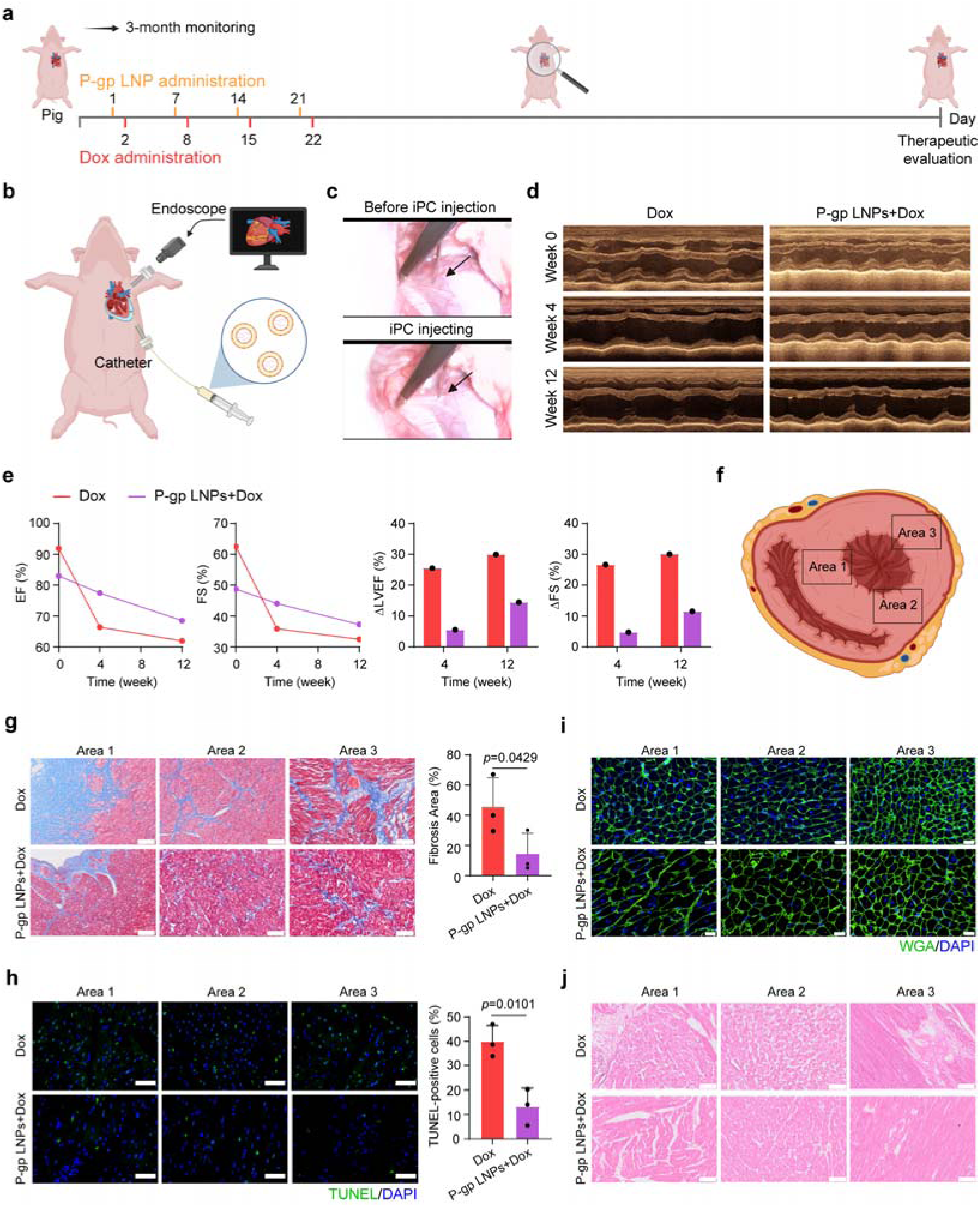
P-gp reprogramming with cardiac drug resistance alleviated Dox-induced cardiotoxicity in pigs. **a**, Schematic illustration of the cardiac P-gp reprogramming by intramyocardial injection of P-gp LNPs and therapeutic evaluation of the cardiac function of Dox-challenged pig. **b**, Schematic illustration of the video-assisted thoracoscopic intrapericardial (iPC) injection of P-gp LNPs in pigs. **c**, Representative snapshots indicating the process of iPC injection. The images were obtained from the thoracoscopy video (Supplementary Movie 1). The injection needle was shown by a black arrow. **d,e**, Representative M-mode echocardiographic images (**d**) and statistical analysis of EF, FS, ΔLVEF, and ΔFS (**e**) in pig with indicated treatments (Dox and P-gp LNPs+Dox) (*n* = 1 independent biological pig per group). **f**, Schematic illustration of heart tissue section for staining. **g**, Representative Masson’s trichrome stained images and statistical analysis of the fibrosis area in the cardiac tissues in pig with indicated treatments. Scale bar, 100 μm. **h**, Representative TUNEL (green) staining images and quantitative analysis of the TUNEL-positive cells of the cardiac tissues in pig with indicated treatments. DAPI staining was employed to visualize the nucleus (blue). Scale bar, 50 μm. **i**, Representative WGA (green) staining images of cardiac tissues in pig with indicated treatments. DAPI staining was employed to visualize the nucleus (blue). Scale bar, 20 μm. **j**, Representative H&E staining images of cardiac tissues in pig with indicated treatments. Scale bar, 100 μm. Significant differences were determined by two-tailed unpaired Student’s *t*-test (**g,h**). Results are shown as meanL±Ls.d. Schematic in **a,b,f** and pig cartoon were created with BioRender.com.

## Discussion

Chemotherapy is a widely employed therapeutic modality in the treatment of a variety of cancers, exhibiting notable therapeutic efficacy. However, observations have highlighted emerging concerns regarding its clinical applications, particularly in view of the associated cardiotoxic adverse effects(48). In particular, doxorubicin (Dox) has been demonstrated to induce a significant dose-dependent cardiotoxicity and subsequent severe pathological cardiomyopathy(43). It was proposed that the lifetime cumulative dose of Dox in humans should not exceed 450-550 mg/m^2^(49). Of note, the dose-dependent cardiotoxicity of Dox is in contradiction with the requirement of multiple scheduled doses of Dox in the chemotherapy of cancer patients. Therefore, the development of a therapeutic approach that can mitigate Dox-induced cardiotoxicity without affecting the anti-cancer efficacy represents a significant challenge in the cancer treatment.

Previous research has illuminated that the cardiotoxicity induced by Dox is mediated by the ROS generation and subsequent iron accumulation, which contribute to cell death in the cardiomyocytes(50). Thus, some studies focused on the depletion the ROS or the modulation of iron metabolism to alleviate the Dox-induced cardiotoxicity, using either small molecules, proteins or nanomedicines(8, 51, 52). Despite the proven efficacy of these strategies, they are constrained by their interaction with downstream pathways subsequent to Dox administration, as opposed to the direct reduction of Dox accumulation in the heart. Moreover, the lack of detailed characterization of the biosafety properties of these novel drugs may result in unknown adverse effects on the body. Our team recently proposed a strategy to alleviate the toxicity of Dox to normal organs by using decoy exosomes, which have been grafted with GC-abundant DNA nanostructures for the capture and detoxification of Dox(53). However, this approach is constrained by the short half-life following systemic administration and the unfavorable distribution patterns.

In addition to Dox-induced cardiotoxicity, the other major obstacle to Dox therapy is the cancer drug resistance. Briefly, the overexpression of transmembrane proteins, such as P-gp, promotes Dox efflux from the cell, resulting in decreased intracellular accumulation of Dox and inhibition of cancer cell apoptosis(54). Our TopCare strategy is dedicated to develop a cardiac drug resistance by mimicking the cancer drug resistance (Fig. 1a). The cardiac drug resistance could be established administration of LNPs encapsulating P-gp mRNAs (P-gp LNPs). The treatment of P-gp LNPs with mice H9c2 and pig PED cells demonstrated the transient overexpression of P-gp proteins in cardiomyocytes, which subsequently resulted in a reduction of intracellular Dox levels and an elevation in cell viability, confirming the in vitro P-gp reprogramming. Additionally, transient overexpression of cardiac P-gp was observed following intramyocardial administration of P-gp LNPs. The production of mRNA is a more straightforward and cost-effective process than that of recombinant proteins or viral vectors(55). The transient reprogramming of cardiomyocytes via mRNA enables precise dosing for the treatment of cardiotoxicity in Dox therapy, while simultaneously reducing the adverse effects of long-term existing therapeutic proteins(38). Of note, the development of cardiac drug resistance did not impede the anticancer efficacy of Dox therapy, as no mRNAs were delivered into tumor tissues.

The biosafety and clinical potential of LNP-mRNAs as therapeutics in the generation of proteins has been demonstrated by the remarkable efficacy of mRNA vaccines in combating the 2019 coronavirus pandemic (59–61). Moreover, clinical trials have been conducted to test the effects of mRNA-based therapeutics in the treatment of cancer, Epstein–Barr virus, and HIV(56). The delivery of therapeutics to the heart with high efficiency has been reported to be challenging and costly(47), especially in large animals. We employed a mini-invasive and highly translatable intrapericardial (iPC) injection technique for P-gp LNPs delivery. The iPC administration of P-gp LNPs demonstrated notable cardioprotective effects against Dox-induced toxicity in pig models.

There are constraints in our present study. The four-component LNPs were prepared in accordance with the methodology outlined in a previous report(57). It would be advantageous to develop novel types of LNPs that are more specific for cardiac applications by modifying the formula of the components in the LNPs. Moreover, although the iPC injection method has recently gained increased interest and has been employed in the cardiac delivery of various therapeutics(58), further investigation into the biosafety and feasibility issues is necessary before its actual clinical applicability can be determined.

In summary, we have provided evidence that the establishment of cardiac drug resistance via P-gp LNPs can protect the heart from the cardiotoxicity in Dox-challenged mice and pig models. The cardiac reprogramming resulted in the overexpression of P-gp proteins and a reduction of Dox levels in cardiomyocytes. The administration of P-gp LNPs to the heart was not only effective in improving cardiac function and attenuating cardiac fibrosis but also did not inhibit the anti-cancer efficacy of Dox therapy. The TopCare strategy presents a potential therapeutic approach for cardio-protection against the Dox-induced cardiotoxicity.

## Methods

### RNA synthesis and encapsulation in lipid nanoparticles

The P-gp mRNA was prepared and synthesized by APE×BIO (Houston, USA). Cloning and endotoxin-free plasmid preparation services were provided by APE×BIO (Houston, USA). The Hyper Scribe T7 kit (APE×BIO, K1409) was employed to generate IVT mRNA from a linearized DNA template containing a 120 nucleotide-long poly(A) tail through in vitro transcription based on T7 polymerase. The nucleotides used were ATP, CTP, GTP, and N1-Methylpseudo-UTP (APE×BIO, B8049). The in vitro transcribed mRNA was then capped and tailed using EZ Cap Reagent AG (3’ OMe) (APE×BIO, B8178). The mRNA was purified using the RNA Clean and Concentrator Kit (APE×BIO, K1069). Subsequently, the mRNA was analyzed by gel electrophoresis (1.5 % agarose) and then kept at −80°C. The lipids were prepared in a molar ratio of SM-102 (APE×BIO, C1042):DMG-PEG 2000 (MCE, HY-112764):DSPC (MCE, HY-W040193):Cholesterol (Aladdin, C104036) = 50:1.5:10:38.5 and were diluted in 100% ethanol. The lipid mixture was then rapidly combined with the P-gp mRNA-containing citrate buffer solution (pH 4.0). The volume ratio of ethanol and the aqueous solution was 1:3. The particle size of P-gp LNPs was characterized using a NanoBrook Omni (Brookhaven Instruments, USA Brookhaven) by dynamic light scattering. The encapsulation rate of mRNA in the P-gp LNP was analyzed via a Quant-iT™ RiboGreen RNA Reagent Kit (Invitrogen Thermo, R11490). The LNPs used in this study are proprietary to Moderna.

### Cell culture and transfection

The rat heart myoblast H9c2 cell line was obtained from the Chinese Academy of Sciences Cell Bank. The cells were then cultured in Dulbecco’s modified Eagle’s medium (DMEM, Gibco) containing fetal bovine serum (10 %, FBS, Biological Industries) and penicillin-streptomycin (2 %, PS, Gibco). P-gp LNPs containing 5 μg P-gp mRNA were added to 1 × 10^6^ H9c2 cells for further experiments.

### Flow cytometry

To verify the overexpression of P-gp and the attenuation of P-gp, H9c2 cells or PED cells were plated into a six-well plate and cultured overnight. P-gp LNPs were then added into the fresh culture medium for transfection. At the appropriate time, the attached H9c2 cells were trypsinized and collected. The cells in the control and transfection groups were then added to sterile PBS. Anti-MDR1/P-glycoprotein antibody (Bioss, bs-1468R) was added, incubated for 1 h at 4°C, and the cells were then washed three times with sterile PBS. The H9c2 cells were resuspended in PBS buffer, stained by Alexa Fluor® 488 (Abcam, ab150077) for 30 min at 4°C in the dark. The cells were then washed three times with sterile PBS. The cells were analyzed using a NovoCyte Quanteon flow cytometer (Agilent, USA). To verify the efflux of Dox, P-gp LNPs were incubated with H9c2 or PED cells for 24 h, after which 1 μM Dox was added. Following an incubation of 24 h, the adherent cells were then trypsinized and collected for subsequent analysis using a NovoCyte Quanteon flow cytometer (Agilent, USA).

### Animal studies

All experiments involving mouse studies were approved and filed by the Ethics Committee at the Tenth Affiliated Hospital of Southern Medical University (IACUC-AWEC-202302016). All experiments involving pig studies were approved by the Guangzhou Huateng Biomedical Technology Co., LTD Ethics Committee (B202403-10). C57BL/6 mice (8-10 weeks, male) were obtained from the Southern Medical University. All the Chinese Bama minipigs were purchased from Guangdong Bright Pearl Biotechnology Co., LTD. The animal experiments were performed strictly following the Regulations for the Administration of Affairs Concerning Experimental Animals (P. R. China).

### Dox-induced cardiomyopathy model in mice

C57BL/6 mice (10 weeks old, male) were anaesthetized and intubated, and were subsequently maintained on a ventilator. P-gp LNPs (10 μg mRNA /20 g) were then injected into the left ventricular myocardium using an insulin needle (0.25×8 mm, Kindly) after the chest was opened. A mouse cardiomyopathy model of Dox-induced cardiomyopathy was established one day later via intraperitoneal injection of doxorubicin hydrochloride (15 mg/kg, MedChemExpress, HY-15142). On day 30, the mice were euthanized after echocardiography. The heart weight and tibia length were then measured. The main organ tissues, including heart, liver, spleen, lung and kidneys were collected for immunostaining.

### Dox-induced cardiomyopathy model in tumor-bearing mice

A B16F10 tumor mouse model was developed via the subcutaneous injection of 1 × 10L B16F10 cells on left side of the C57BL/6 mice (8 weeks, male). When the tumor grew to approximately 50 mm³ in volume, mice were anaesthetized and intubated, and then maintained on a ventilator. The mice were subsequently administrated an intramyocardial injection of P-gp LNP (10 μg mRNA /20 g). The Dox (15 mg/kg) was then administered intraperitoneally 24 h later. The tumor volume and mice’s body weight were measured every two days during the course of the treatment. The tumor volume of each mouse was calculated according to the formula *V* (volume) = length × width² /2. On day 14, all mice were then euthanized using COL asphyxiation, followed by dissection. The main organ tissues (heart, spleen, liver, lung, kidneys, and tumor) were harvested for analysis.

### Mouse echocardiography

Following a 30-day administration period, the mice were anaesthetized using 1.5% isoflurane and then placed in a supine position for transthoracic echocardiography (Vevo 3100LT, FUIJIFILM). Morphological and functional parameters were quantified from M-mode/2D images using VevoLAB software (FUIJIFILM VisualSonics).

### Western blot

The total proteins were isolated from the specific cells and tissues at the designated time point. A equal amount of proteins in each sample was then separated using a 7.5% gel prepared by the Color PAGE Gel Rapid Preparation Kit (Epizyme, PG111). The proteins were transferred to PVDF membranes (Millipore, 03010040001). After incubation with 5% BSA for 2 h at room temperature for blocking, the PVDF membranes were incubated with antibodies against P-glycoprotein (Abcam, ab170904, 1:3000) and β-tubulin (Proteintech, 10094-1-AP, 1:3000) overnight at 4°C. After washing, the PVDF membranes were then incubated with HRP-conjugated goat anti-rabbit IgG(H+L) (Proteintech, SA00001-2, 1:3000) for 2 h at room temperature. Then, the PVDF membranes were washed, exposed (Tanon 1600), and the data were analyzed using Image J.

### HPLC Analysis of Dox in the heart

The heart was excised 6 h after intraperitoneal injection of Dox and dried with filter paper, and weighed. The heart was homogenized in 300 μl of acetonitrile, shaken overnight in a four-degree shaker, and the resulting supernatant was then centrifuged. The supernatant was mixed with a known quantity of the internal standard for Dox and then injected. HPLC analyses were performed with a Waters Alliance series instrument. The column utilized was a YMC ODS (particle sizes, 5 μm; pore sizes, 120 Å; 250×4.6mm I.D.; AA12S05-2546WT). The flow rate was set at 1.0 mL/min, and the column temperature was maintained at 40°C. The mobile phase was composed of a water/acetonitrile (68:32) solution containing trifluoroacetic acid (0.1%). The chromatograms were determined by recording the UV absorption (λ = 254 nm).

### In vivo biodistribution

Luc LNPs were administrated into mice via intramyocardial injection, and the D-luciferin (150 mg/kg) were intraperitoneally injected at 24 h. The mice were then anesthetized with isoflurane 10 min later, and subjected to whole-body imaging using an IVIS Spectrum (PerkinElmer). After that, the mice were sacrificed, and main organ tissues (heart, spleen, liver, lung, kidneys, and tumor) were then collected for subsequent ex vivo imaging. The imaging data were quantified and analyzed using Living Image software (PerkinElmer).

### Histological analysis

The tissues were firstly fixed in 4% paraformaldehyde (PFA), and then immersed in 30% sucrose solution. Then tissues were ready for use when sank to the bottom in the solution. The tissues were embedded in OCT and 10-μm tissue sections were cut using a cryostat (Leica CM3050 S). The tissue sections were incubated with 0.1% Triton X-100 solution at room temperature for 10 min and then incubated with 5% BSA at room temperature for 2 h. Heart sections injected with eGFP LNPs were incubated with α-sarcomeric actinin antibody (1:500, Abcam, ab317695), GFP antibody (1:500, Proteintech, 66002-1-Ig) and α-smooth muscle actin antibody (1:500, Proteintech, 14395-1-AP) overnight at 4°C. After washing three times with sterile PBS, the tissue sections were stained with Alexa Fluor® 488 antibody (Abcam, ab150113, 1:500) or Alexa Fluor® 647 antibody (Abcam, ab150079, 1:500) at room temperature for 2 h and then washed with PBS. The sections of main organ tissues from mice injected with Luc LNPs were incubated with anti-firefly luciferase antibody (1:500, Abcam, ab185924) overnight at 4°C. After washing three times with sterile PBS, the tissue sections were stained with Alexa Fluor® 488 antibody (Abcam, ab150077) at room temperature for 2 h and then washed with sterile PBS. The sections were then stained with DAPI. TUNEL staining of the tissue sections was conducted with a One-Step TUNEL Apoptosis Assay Kit (Beyotime, C1086). Tissue morphology was evaluated through the use of a Haematoxylin and Eosin (H&E) staining kit (Beyotime, C0105S) on sections of main organ tissues. The sections of heart were subjected to staining using the Masson’s trichrome staining kit (Solarbio, G1340). Selective labelling of plasma membranes was achieved using the wheat germ agglutinin (WGA, Thermofisher, W11261). The distinction between collagen and muscle fibers was made on the basis of the size of the anionic dye molecules and the varying degrees of tissue penetration.

### Pig studies

The efficacy of P-gp LNPs was initially validated at the cellular level with PED (porcine endothelial cell, procell, CL-0482) cell line. Subsequently, P-gp LNPs comprising 150 μg of P-gp mRNA were administered into pigs via iPC injection on a weekly basis for a period of four weeks into male Bama minipigs (approximately 25 kg), in accordance with the methodology previously established. Dox (1 mg/kg) was then intravenously administered into the pigs via the ear margin 24 h later on a weekly basis for a period of four weeks. Echocardiography and electrocardiography were conducted at different time points (day 0, week 4 and week 12 post intervention) to to evaluate the cardiac function. At week 12, the pigs were sacrificed via euthanasia, and the heart was excised for histological analysis.

### Statistical analysis

All the experiments were conducted a minimum of three times. The data were analyzed using the GraphPad Prism 8 (version 8.0.2.263), and statistical significance was assessed by one-way ANOVA and Bonferroni’s multiple comparisons test, two-way ANOVA and Bonferroni’s multiple comparisons test, or two-tailed unpaired Student’s *t*-test. Results are shown as meanL±Ls.d.

## Data availability

All data supporting the conclusions of this study are presented in the article and the Supplementary Information.

## Acknowledgements

This work was supported by grants from the National Key R&D Program of China (2022YFB3808300 to Z.L.), National Natural Science Foundation of China (32322045 to Z.L., 32301162 to F.W., 22207050 to L. Li, 82202339 to X.W.). The schemes in figures were created with Biorender.com.

## CRediT authorship contribution statement

Z.L., F.W., and H.L. conceived the study and designed the experiments. Y. Zhang, and W.L. led and contributed to all the experiments together. Y.X. and X.W. contributed to the preparation and characterization of P-gp LNPs. S.M., L. Li, J.D. contributed to the cell experiments. L. Luo, J.A., Q.L., and Y. Zeng contributed to the animal experiments. Y. Zhang, W.L., and F.W. analyzed the data and prepared the figures. Y. Zhang and W.L. wrote the paper, and Y. Zhang, W.L., H.L., F.W. and Z.L. reviewed and corrected the paper with contributions from all authors.

## Additional information

**Supplementary information** The online version contains supplementary material available at

