## Supplemental Information for "Cardiac Reprogramming with Drug Resistance Alleviates Doxorubicin-induced Cardiotoxicity in Mice and Pigs"

### SUPPLEMENTARY FIGURES

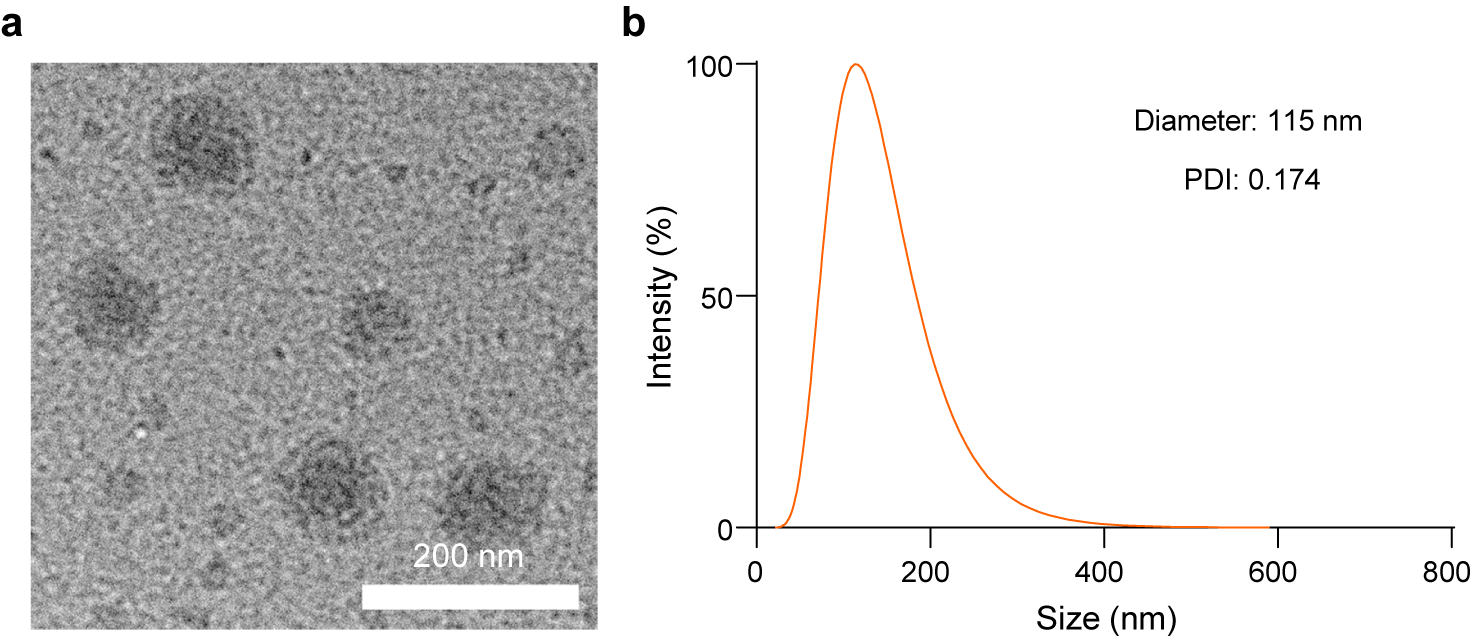

Supplementary figure S1**. Characterizations of P-gp LNPs. a**, Representative transmission electron microscopy (TEM) image of the P-gp LNPs. Scale bar, 200 nm. Experiments were replicated in triplicate. **b**, Size distribution of P-gp LNPs determined by dynamic light scattering. Experiments were replicated in triplicate.

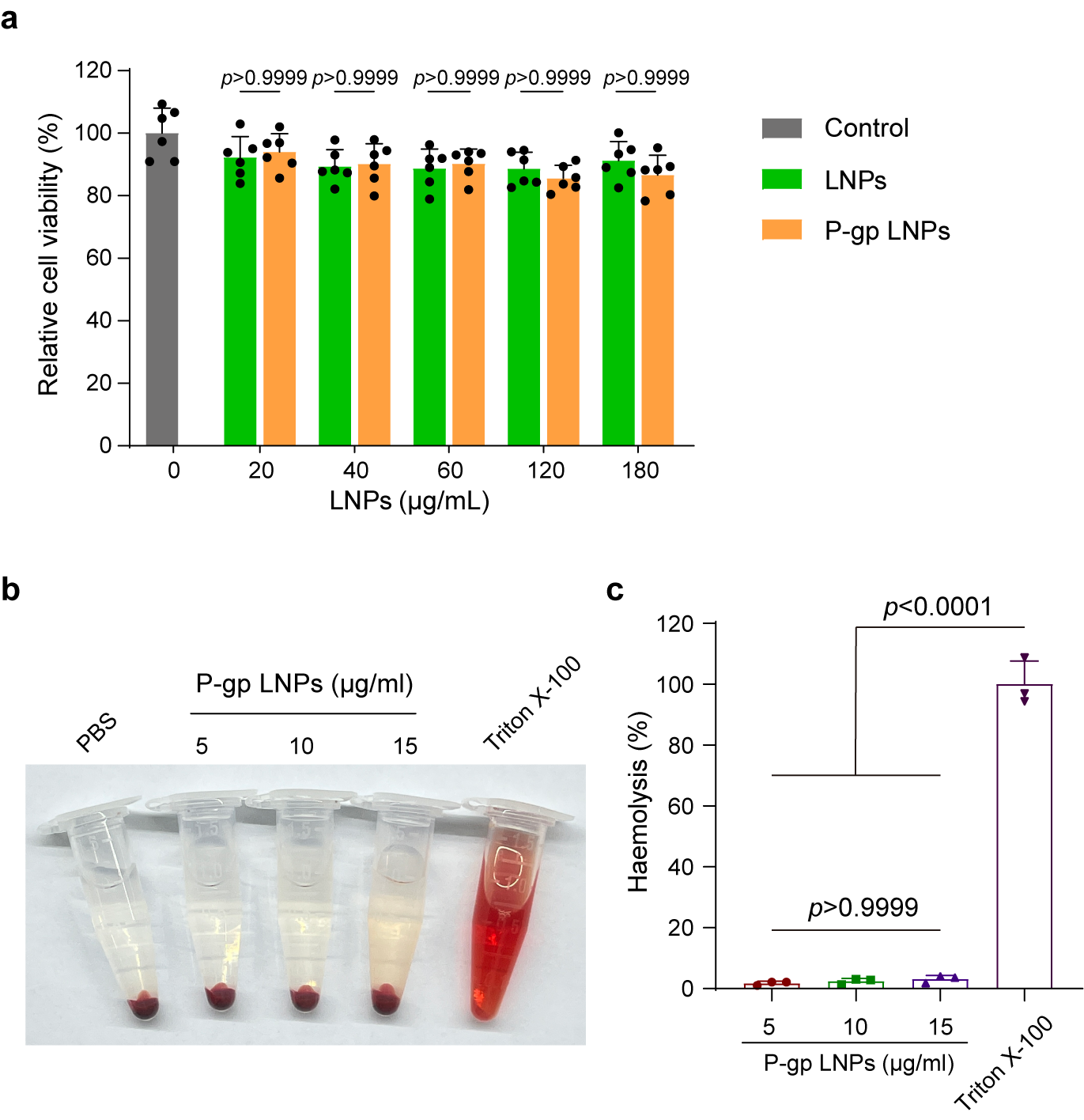

Supplementary figure S2**. In vitro biosafety of P-gp LNPs. a**, Relative cell viability of H9c2 cells treated with LNPs and P-gp LNPs at different concentrations determined by CCK-8 assays (*n* = 6 independent biological samples). **b,c**, Hemolysis analysis (**b**) and quantitative analysis (**c**) of P-gp LNPs at different concentrations at pH 7.4. Triton X-100 treatment was used as a positive control for hemolysis. Significant differences were assessed by two-way ANOVA and Bonferroni’s multiple comparisons test (**a**). Results are presented as mean ± s.d.

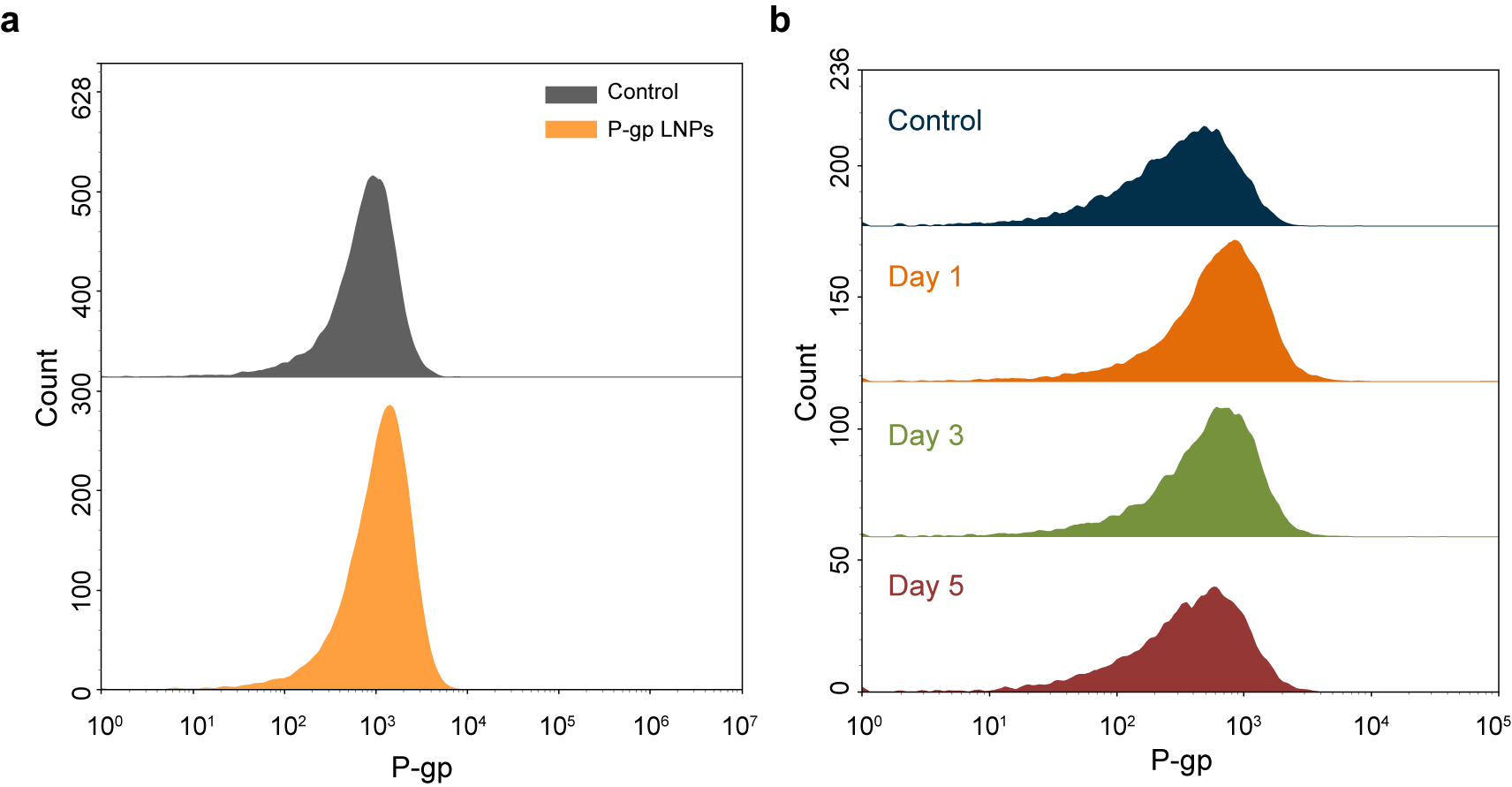

Supplementary figure S3**. Flow cytometry analysis of P-gp expression and attenuation. a**, Flow cytometry analysis of P-gp expression in H9c2 cells 24 h after treated with indicated treatments (Control and P-gp LNPs). Experiments were replicated in triplicate. **b**, Flow cytometry analysis of the attenuation of the overexpressed P-gp in H9c2 cells after incubation with P-gp LNPs over a 5-day period. Experiments were replicated in triplicate.

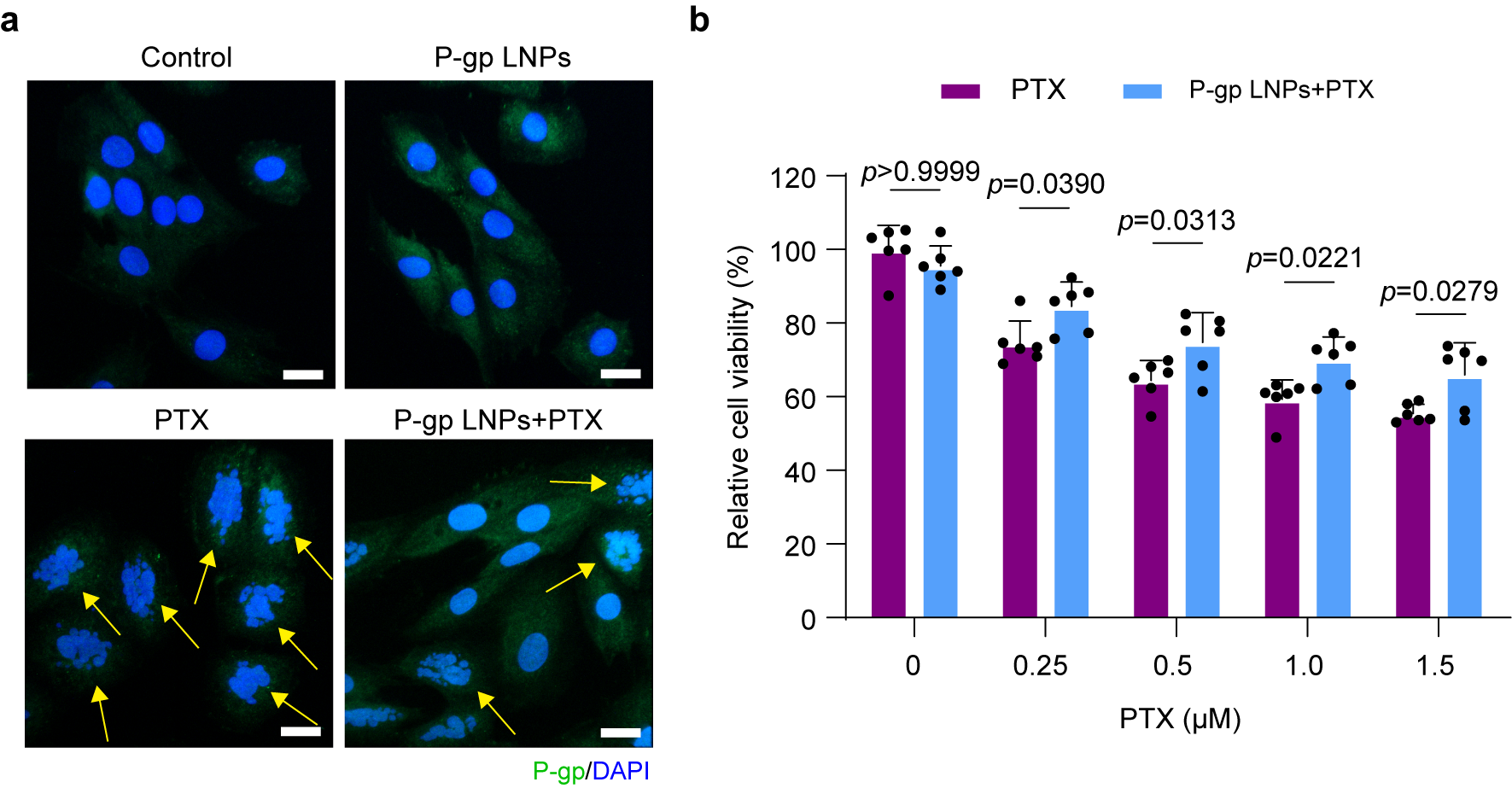

Supplementary figure S4**. In vitro P-gp overexpression reduced the toxicity of paclitaxel (PTX) to cardiomyocytes. a**, Representative immunofluorescence staining images of H9c2 cells with the indicated treatments (PTX and P-gp LNPs+PTX). Cells were treated with a 24 h pretreatment of PBS or P-gp LNPs prior to incubation with PTX for 24 h. P-gp was stained green. DAPI staining was used to show the nucleus (blue). The mitotic spindle defect was shown by a yellow arrow. Scale bar, 20 μm. Experiments were replicated in triplicate. **b**, Relative cell viability of H9c2 cells with indicated treatments (PTX and P-gp LNPs+PTX) determined via CCK-8 assays (*n* = 6 independent biological samples). Significant differences were assessed by two-way ANOVA and Bonferroni’s multiple comparisons test (**b**). Results are presented as mean ± s.d.

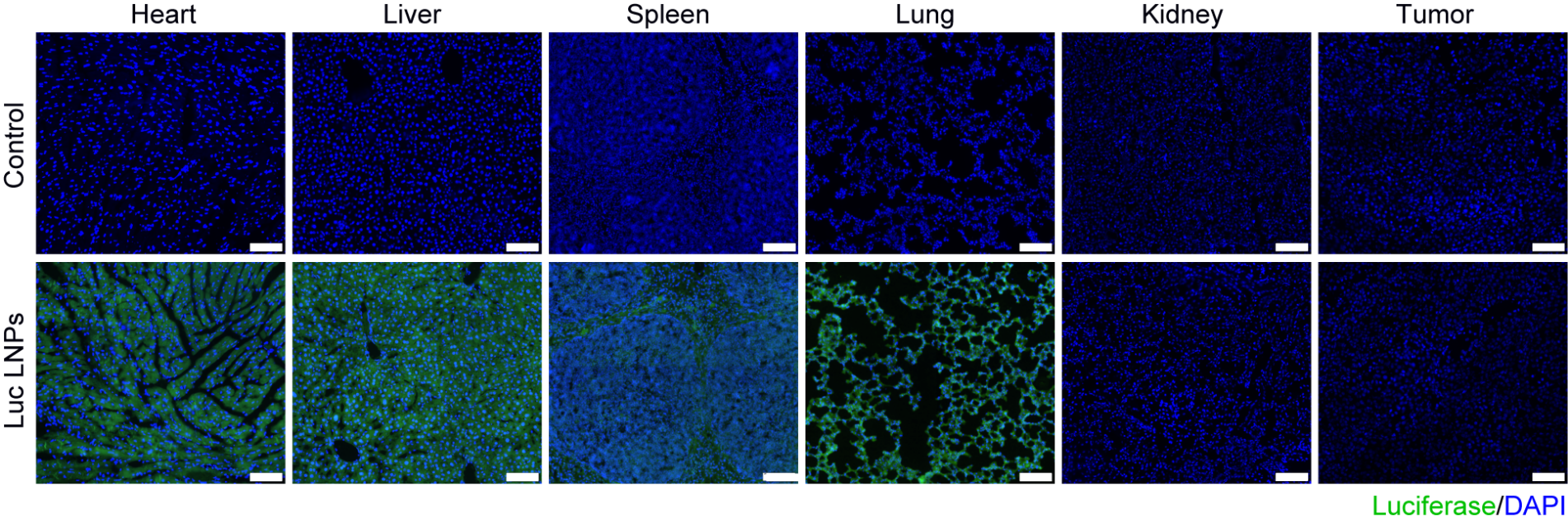

Supplementary figure S5**.** **Immunofluorescence staining of luciferase in major tissues.** Representative immunofluorescence staining images of luciferase (green) expression in major tissues (heart, liver, spleen, lung, kidney, tumor) 24 h after intramyocardial injection of Luc LNPs in mice. DAPI staining was used to show the nucleus (blue). Scale bar, 100 μm. Experiments were replicated in triplicate.

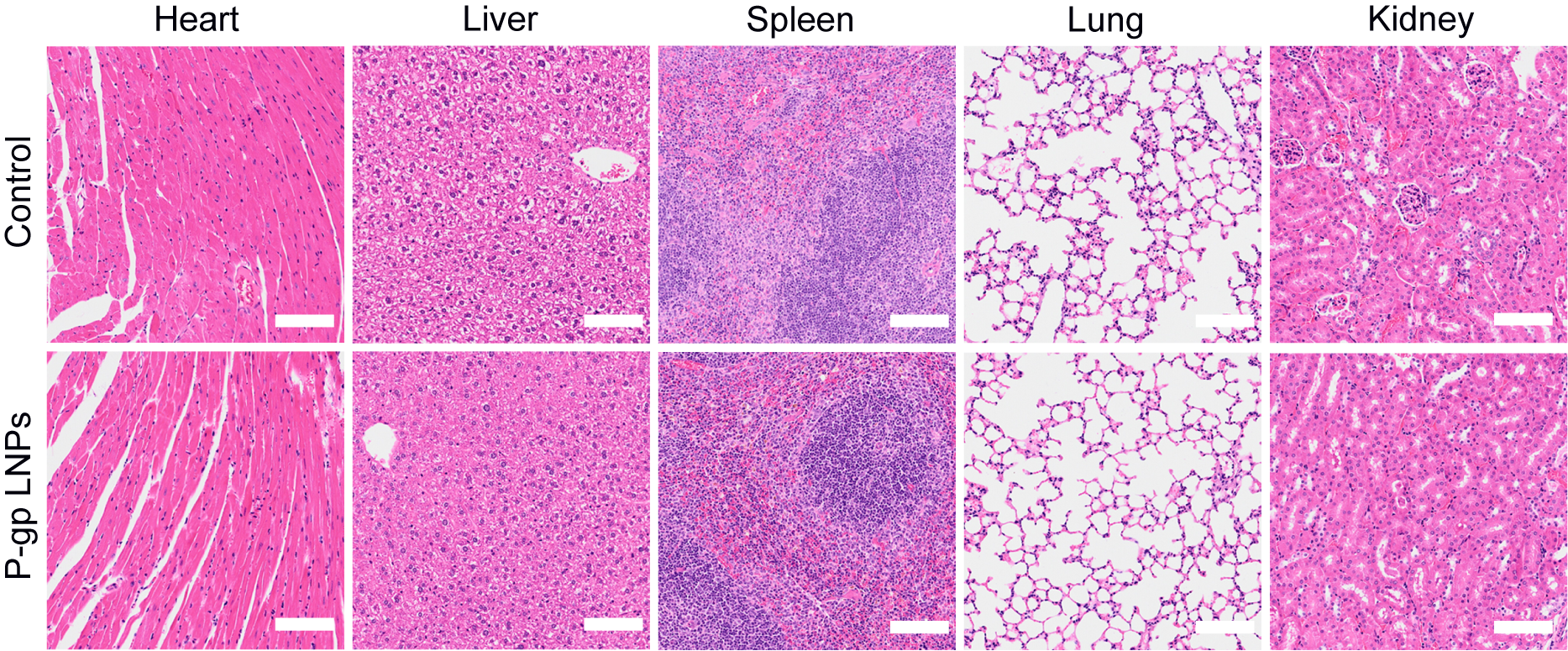

Supplementary figure S6**.** **H&E staining of major tissues.** Representative H&E staining images of major tissues (heart, liver, spleen, lung, kidney) collected on day 7 after intramyocardial injection of P-gp LNPs in mice. Scale bar, 100 μm.

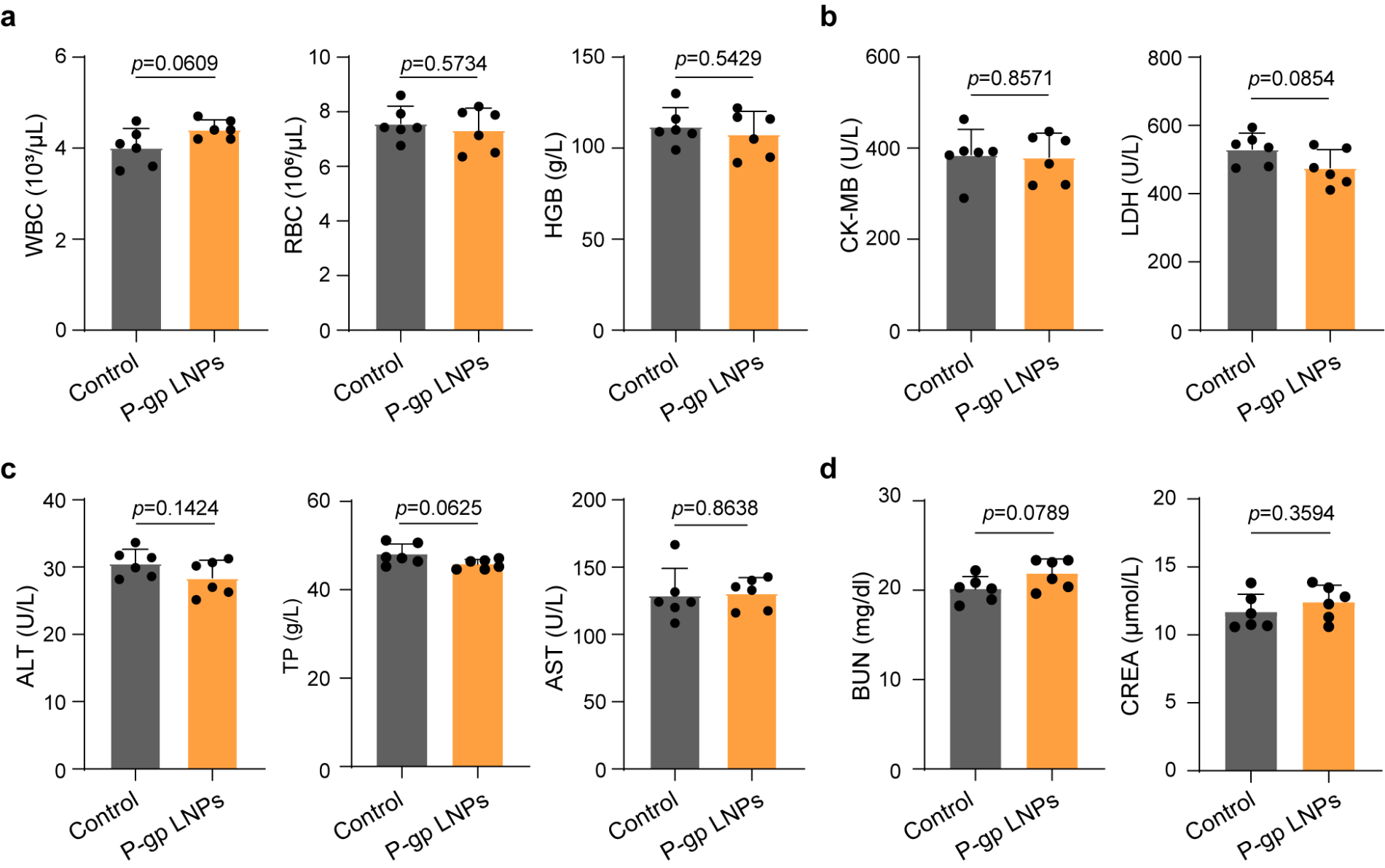

Supplementary figure S7**. Blood routine tests and biochemical analysis.** **a**, Routine blood tests to determine the hematological parameters (*n* = 6 independent biological mice per group). WBC: white blood cells; RBC: red blood cells; HGB: hemoglobin. **b**, Biochemical analysis of the key biomarkers of cardiac toxicity (*n* = 6 independent biological mice per group). CK-MB: creatine kinase−myocardial band; LDH: lactate dehydrogenase. **c**, Biochemical analysis of the key biomarkers of liver toxicity (*n* = 6 independent biological mice per group). ALT: alanine transaminase; TP: total protein; AST: aspartate transaminase. **d**, Biochemical analysis of the key biomarkers of kidney toxicity (*n* = 6 independent biological mice per group). BUN: blood urea nitrogen; CREA: creatinine. Blood samples were collected on day 7 after intramyocardial injection of P-gp LNPs in mice. Significant differences were assessed by two-tailed unpaired Student’s *t*-test (**a-d**). Results are presented as mean ± s.d.

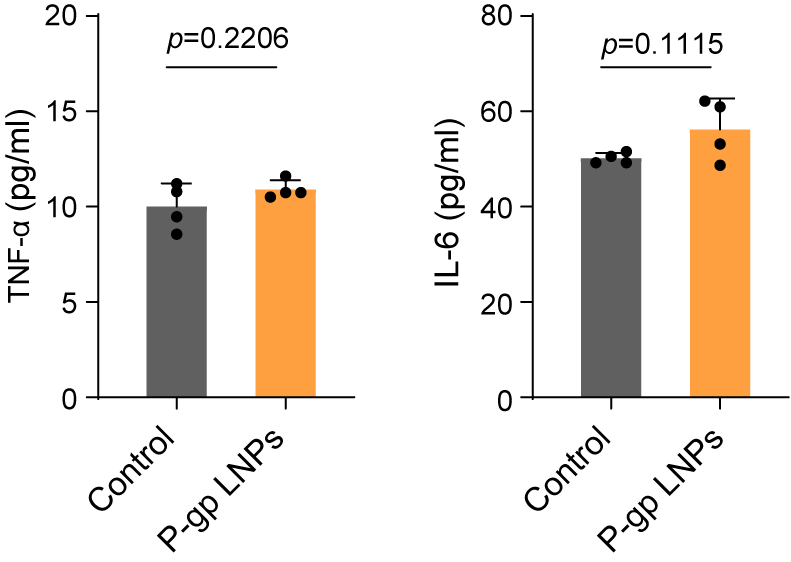

Supplementary figure S8**. ELISA analysis of pro-inflammatory cytokines.** ELISA analysis of the serum pro-inflammatory factors, including TNF-α and IL-6 (*n* = 4 independent biological mice per group). Blood samples were collected 24 h after intramyocardial injection of P-gp LNPs in mice. Significant differences were assessed by two-tailed unpaired Student’s *t*-test. Results are presented as mean ± s.d.

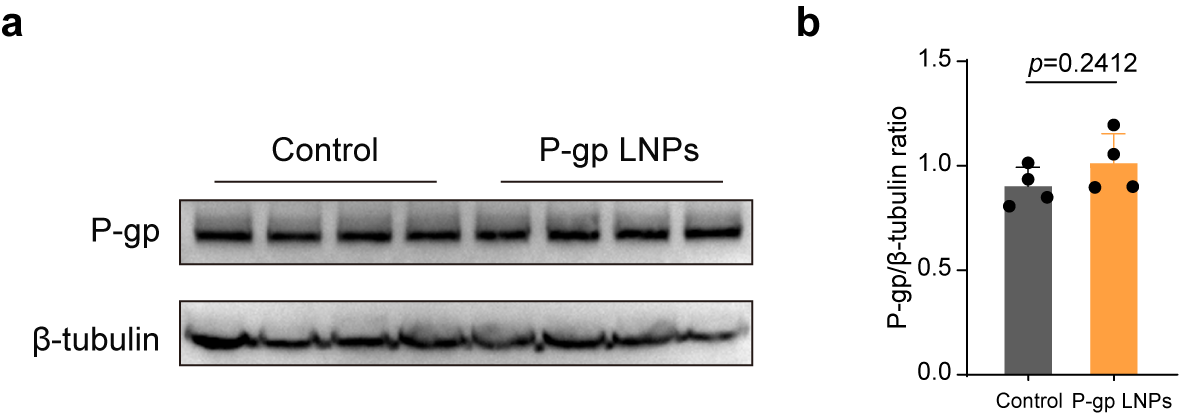

Supplementary figure S9**. P-gp expression in tumor tissues.** **a,b**, Western blotting analysis (**a**) and statistical analysis (**b**) of P-gp expression in the tumors of mice with indicated treatments (Control and P-gp LNPs) for 24 h (*n* = 3 independent biological mice per group). Significant differences were assessed by two-tailed unpaired Student’s *t*-test (**b**). Results are presented as mean ± s.d.

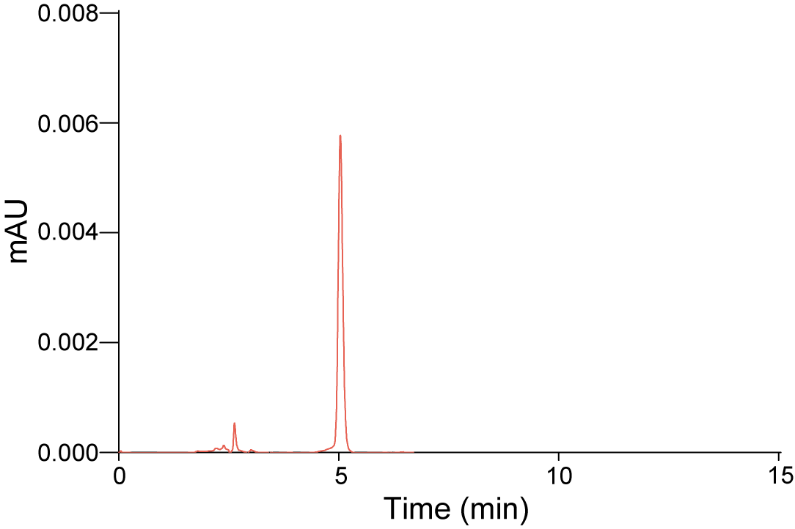

Supplementary figure S10**.** **High-performance liquid chromatography (HPLC) analysis of the doxorubicin hydrochloride.**

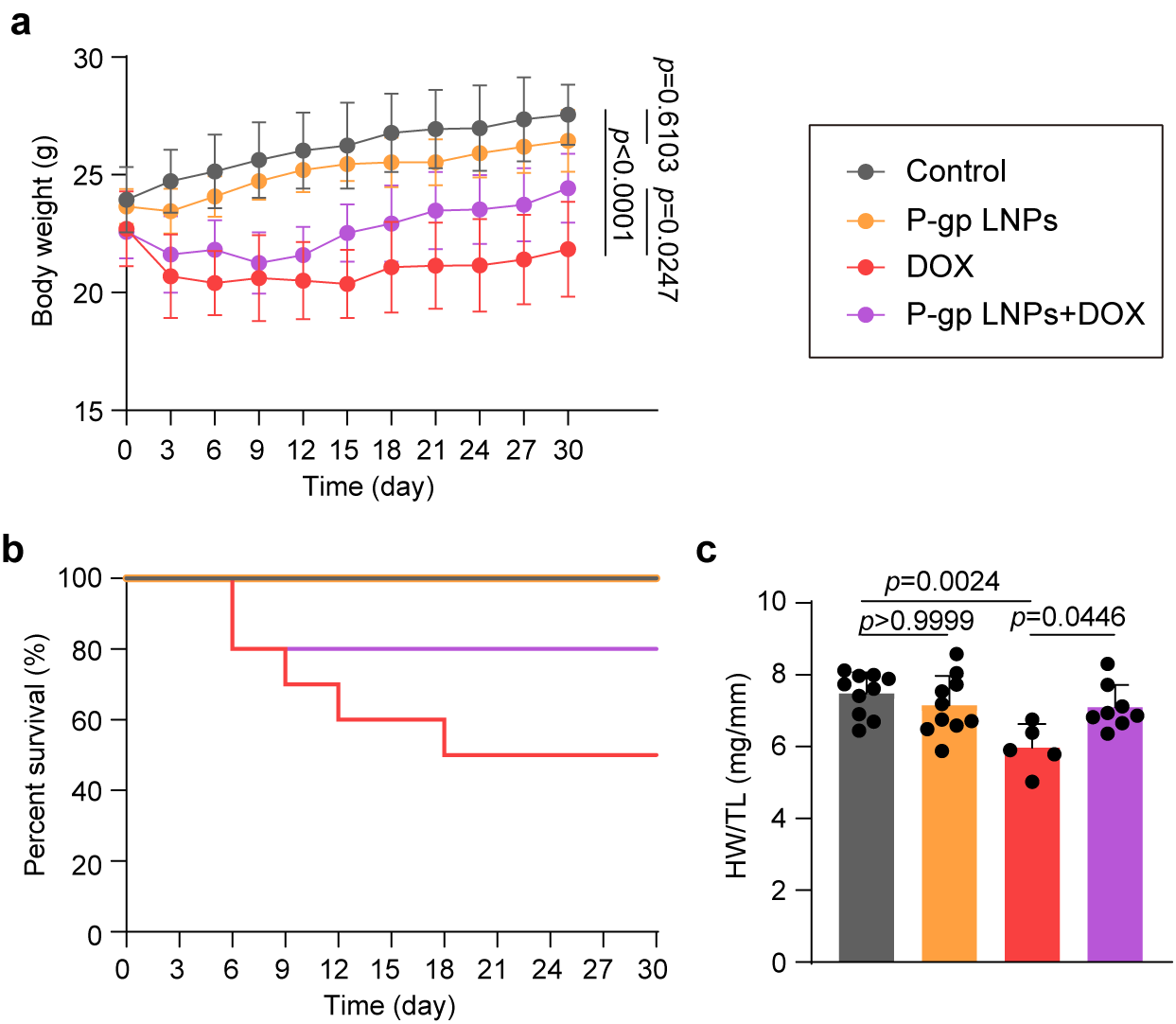

Supplementary figure S11**. P-gp LNPs alleviate Dox-induced cardiotoxicity in mice. a,** Relative body weight changes over time in each group with indicated treatments (Control, P-gp LNPs, Dox, and P-gp LNPs+Dox). (*n* = 10 independent biological mice per group). **b**, Survival curves of mice with indicated treatments (Control, P-gp LNPs, Dox, and P-gp LNPs+Dox) (*n* = 10 independent biological mice per group). **c**, HW/TL of the mouse in each group with different treatments (*n* = 10, 10, 5, and 8 independent biological mice per group, respectively). Significant differences were assessed by one-way ANOVA and Bonferroni’s multiple comparisons test (**c**). Results are presented as mean ± s.d.

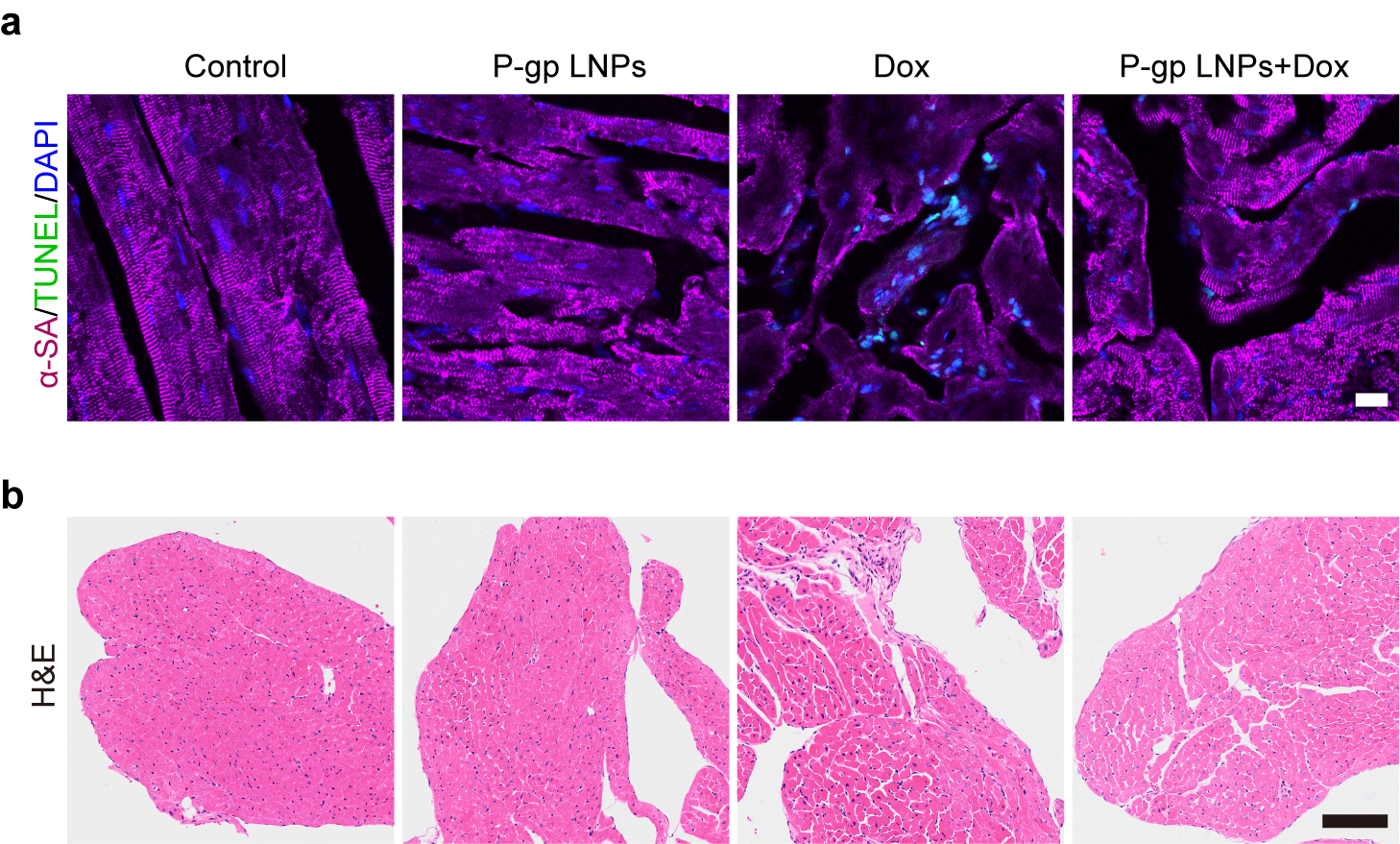

Supplementary figure S12**.** **TUNEL staining and H&E staining of cardiac tissues.** **a**, Representative TUNEL (green) staining images of the cardiac tissues from mice with indicated treatments (Control, P-gp LNPs, Dox, and P-gp LNPs+Dox) (n = 3 biologically independent mice per group). α-SA was stained pink. DAPI staining was used to show the nucleus (blue). Scale bar, 20 μm. **b**, Representative H&E staining images of cardiac tissues from mice with indicated treatments (Control, P-gp LNPs, Dox, and P-gp LNPs+Dox). Scale bar, 100 μm. Experiments were replicated in triplicate.

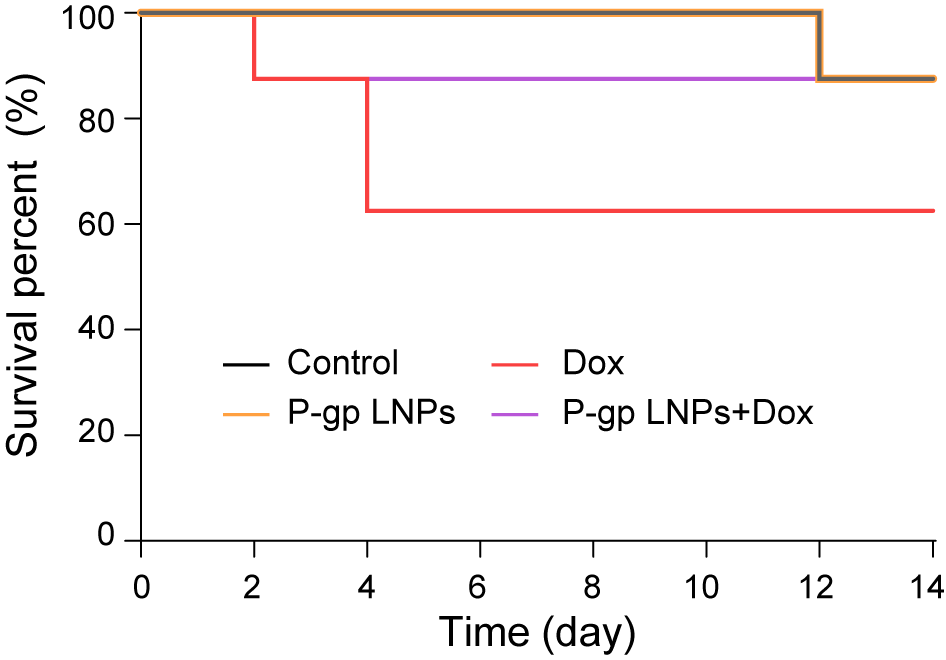

Supplementary figure S13**.** **Survival curves of B16F10 tumor-bearing mice with indicated treatments.** Treatments include: Control, P-gp LNPs, Dox, and P-gp LNPs+Dox) (*n* = 8 independent biological mice per group).

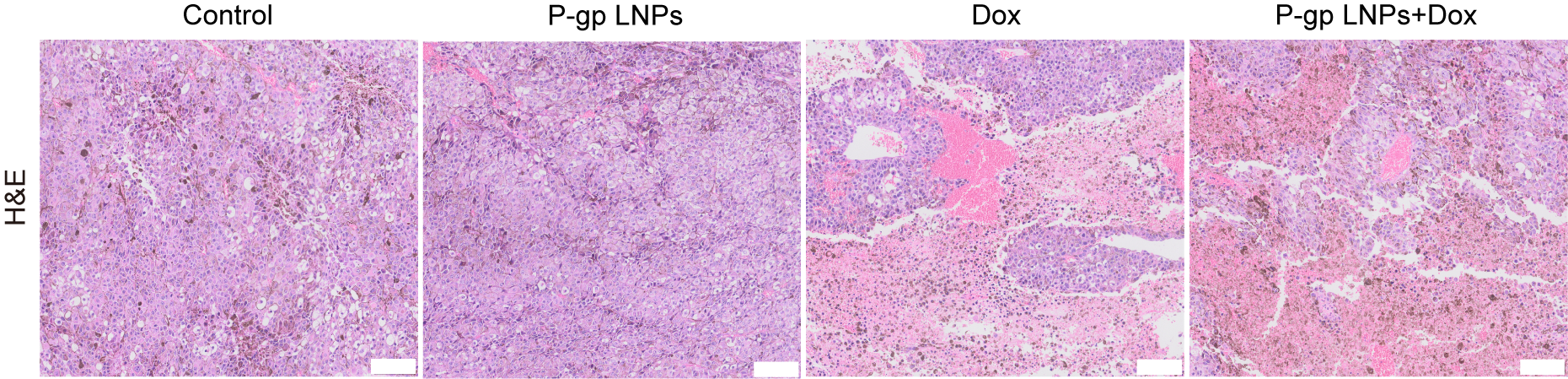

Supplementary figure S14**. H&E staining of tumor tissues.** Representative H&E staining images of tumor tissues dissected from mice with indicated treatments (Control, P-gp LNPs, Dox, and P-gp LNPs+Dox). Scale bar, 100 μm. Experiments were replicated in triplicate.

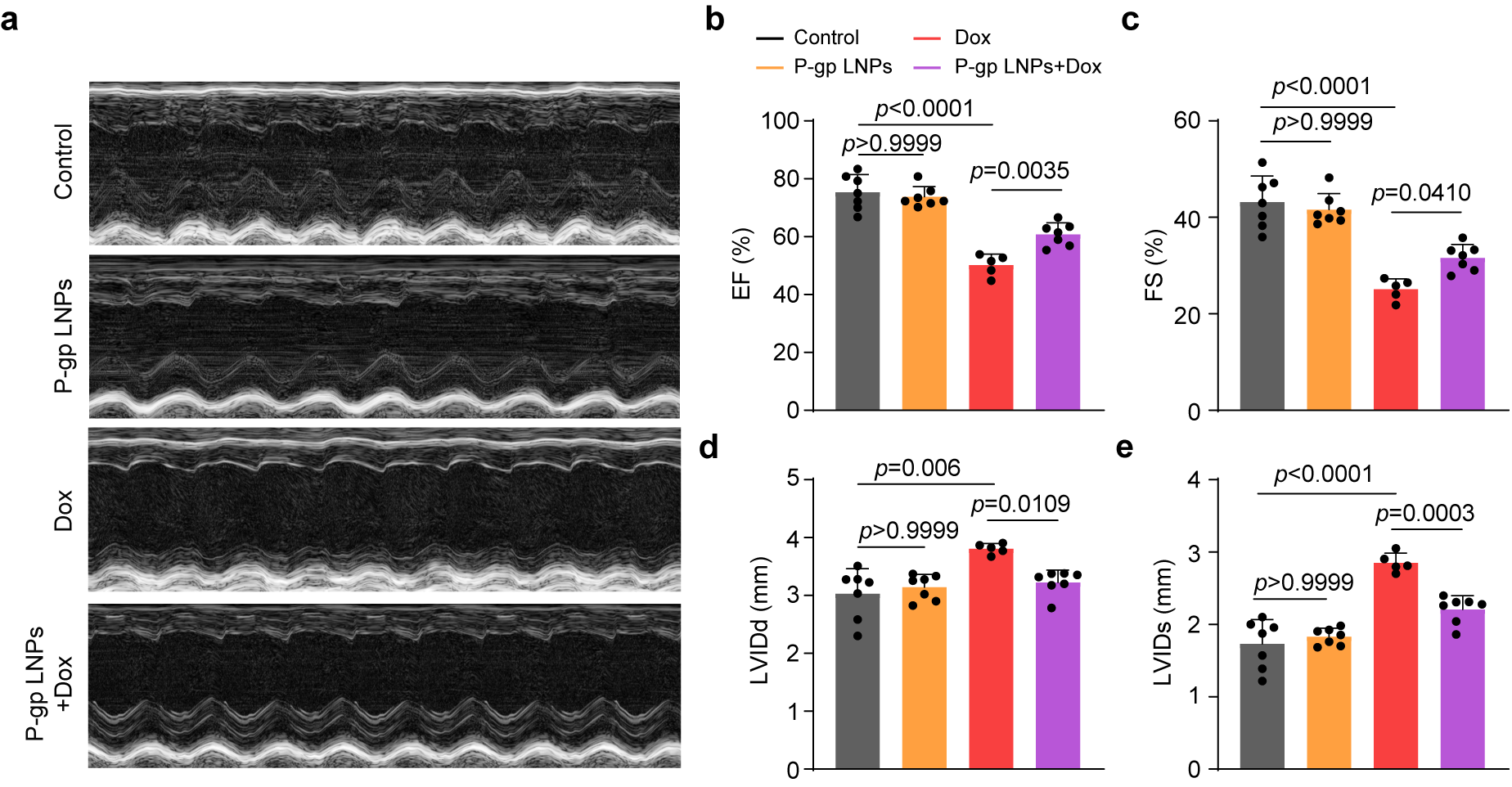

Supplementary figure S15**. In vivo cardioprotective effects in tumor-bearing mice.** **a-e**, Representative M-mode echocardiography images (**a**) and quantitative analysis of EF (**b**), FS (**c**), LVIDd (**d**), and LVIDs (**e**) of mice with indicated treatments (Control, P-gp LNPs, Dox, and P-gp LNPs+Dox) (*n* = 7, 7, 5, and 7 independent biological mice per group, respectively). Significant differences were assessed by one-way ANOVA and Bonferroni’s multiple comparisons test (**b-e**). Results are presented as mean ± s.d.

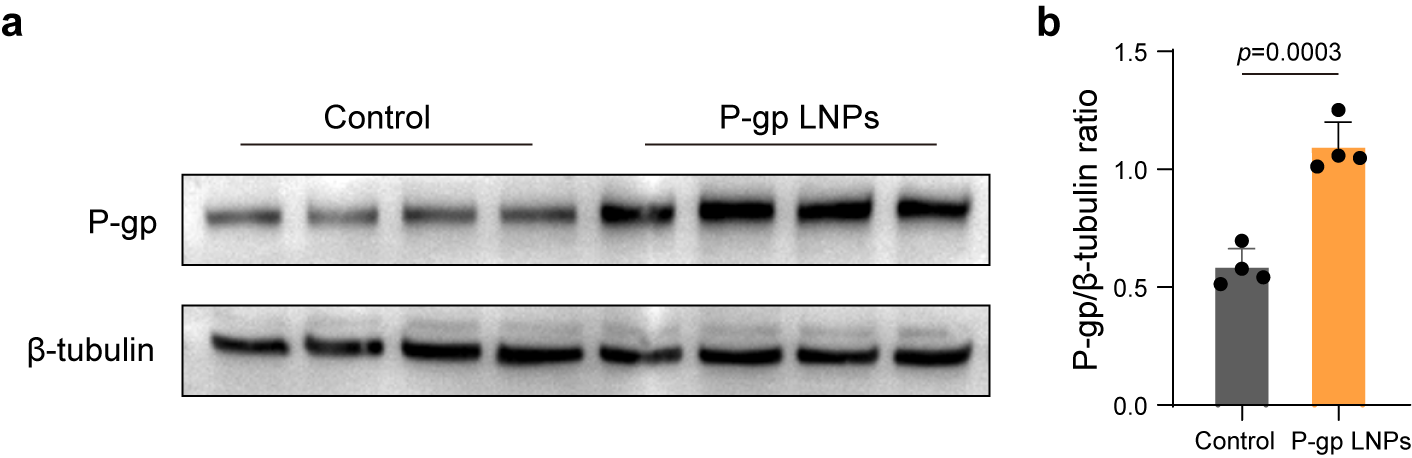

Supplementary figure S16**. P-gp expression in PED cells. a,b**, Western blotting analysis (**a**) and statistical analysis (**b**) of P-gp expression in PED cells treated with PBS (Control) and P-gp LNPs for 24 h (*n* = 4 independent biological samples). Significant differences were assessed by two-tailed unpaired Student’s *t*-test (**b**). Results are presented as mean ± s.d.

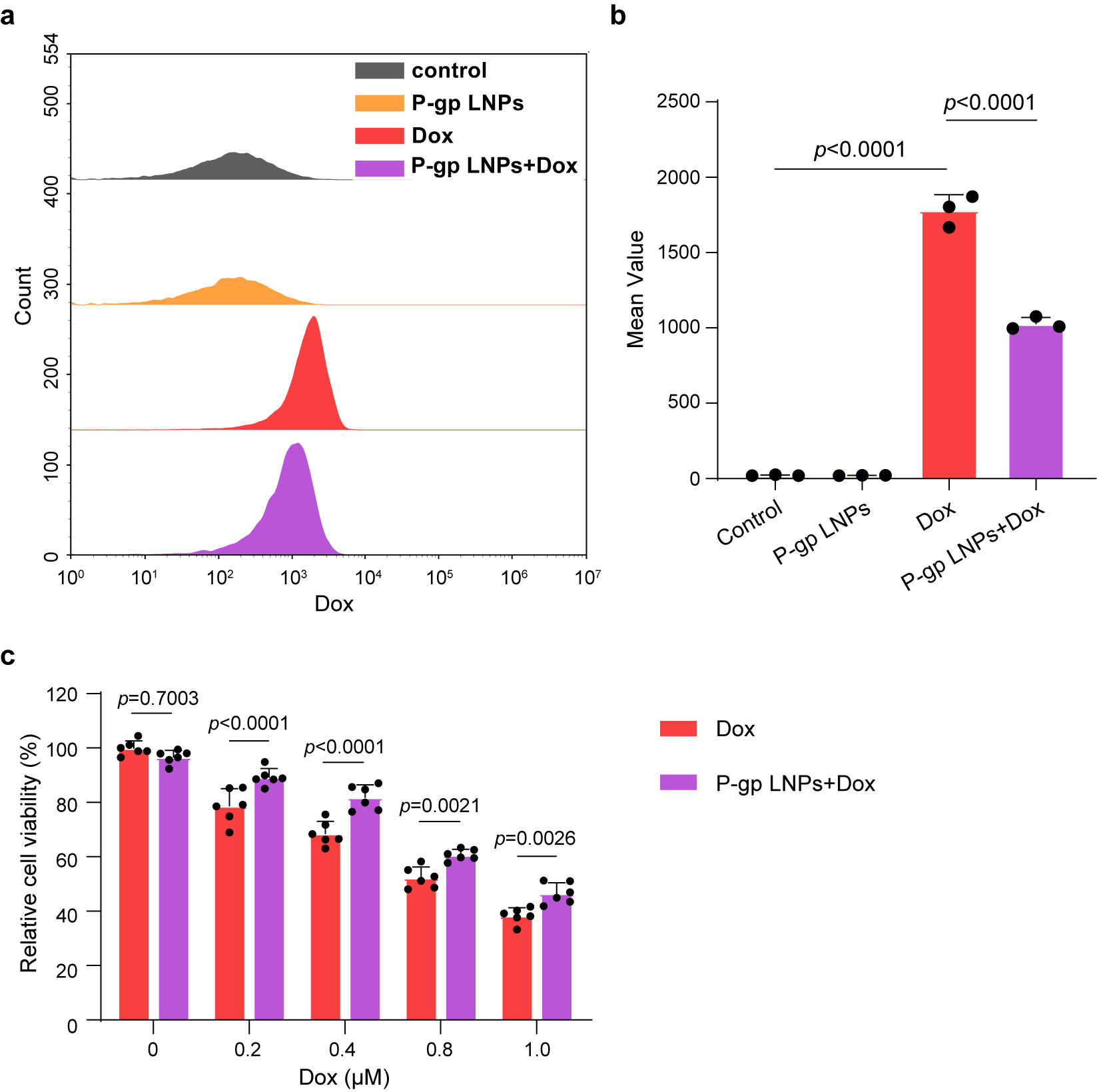

Supplementary figure S17**. P-gp LNPs reduced intracellular Dox in PED cells. a,b**, Flow cytometry analysis (**a**) and quantitative mean value (**b**) of intracellular Dox in PED cells with indicated treatments (Control, P-gp LNPs, Dox, and P-gp LNPs+Dox) (*n* = 3 independent biological samples). **c**, Relative cell viability of PED cells with indicated treatments (Dox and P-gp LNPs+Dox) determined via CCK-8 assays (*n* = 6 independent biological samples). Significant differences were assessed by one-way ANOVA and Bonferroni’s multiple comparisons test (**b**) and two-way ANOVA and Bonferroni’s multiple comparisons test (**c**). Results are presented as mean ± s.d.

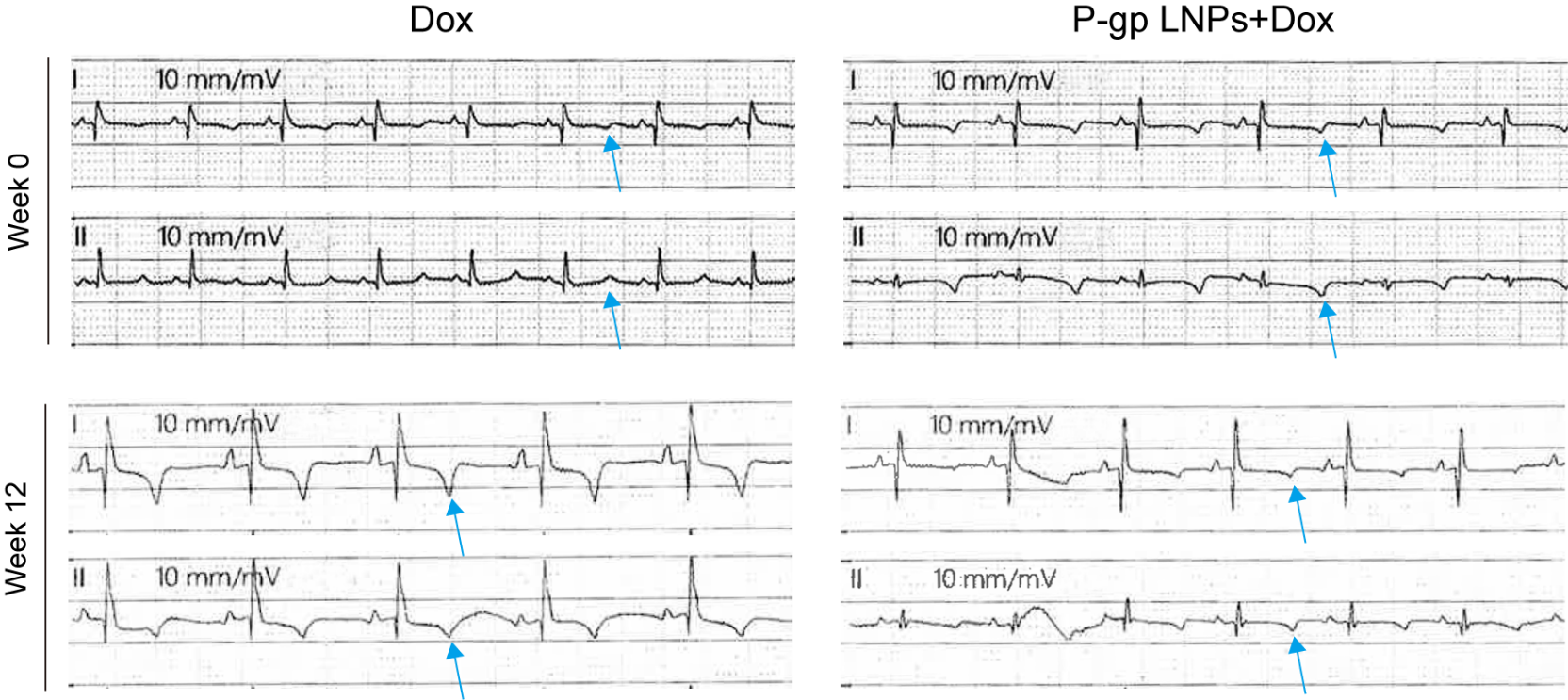

Supplementary figure S18**. Electrocardiogram of pigs.** Representative electrocardiogram images of pigs at week 0 and week 12 after indicated treatments (Dox and P-gp LNPs+Dox). The T-wave was shown by a blue arrow.

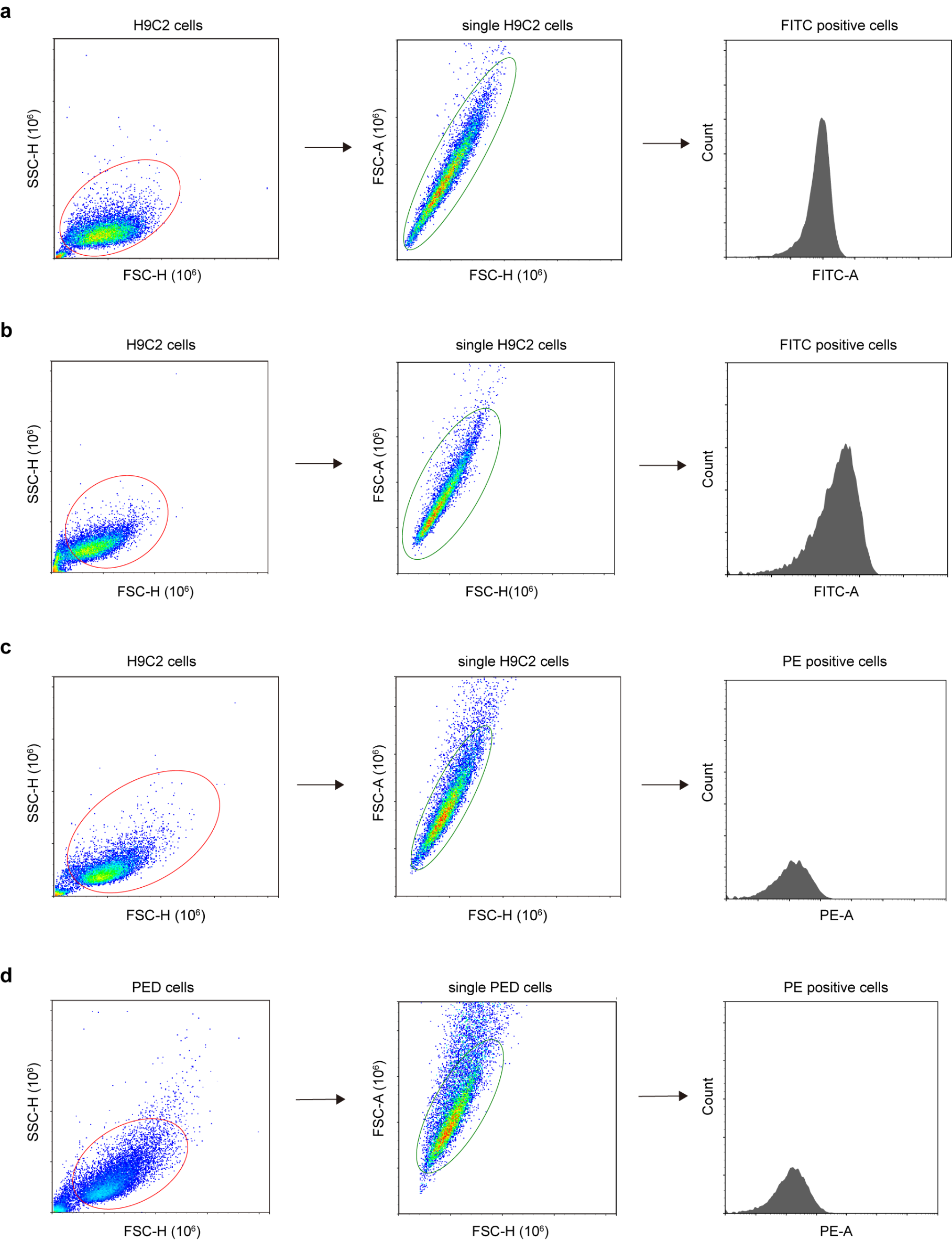

Supplementary figure S19**. Gating strategies in flow cytometry. a,** Gating strategy for flow cytometry analysis of P-gp expression in H9c2 cells with indicated treatments (Control and P-gp LNPs). **b**, Gating strategy for flow cytometry analysis of the attenuation of the overexpressed P-gp in H9c2 cells after incubation with P-gp LNPs. **c**, Gating strategy for flow cytometry analysis of intracellular Dox in H9c2 cells with indicated treatments (Control, P-gp LNPs, Dox, and P-gp LNPs+Dox). **d**, Gating strategy for flow cytometry analysis of intracellular Dox in PED cells with indicated treatments (Control, P-gp LNPs, Dox, and P-gp LNPs+Dox).

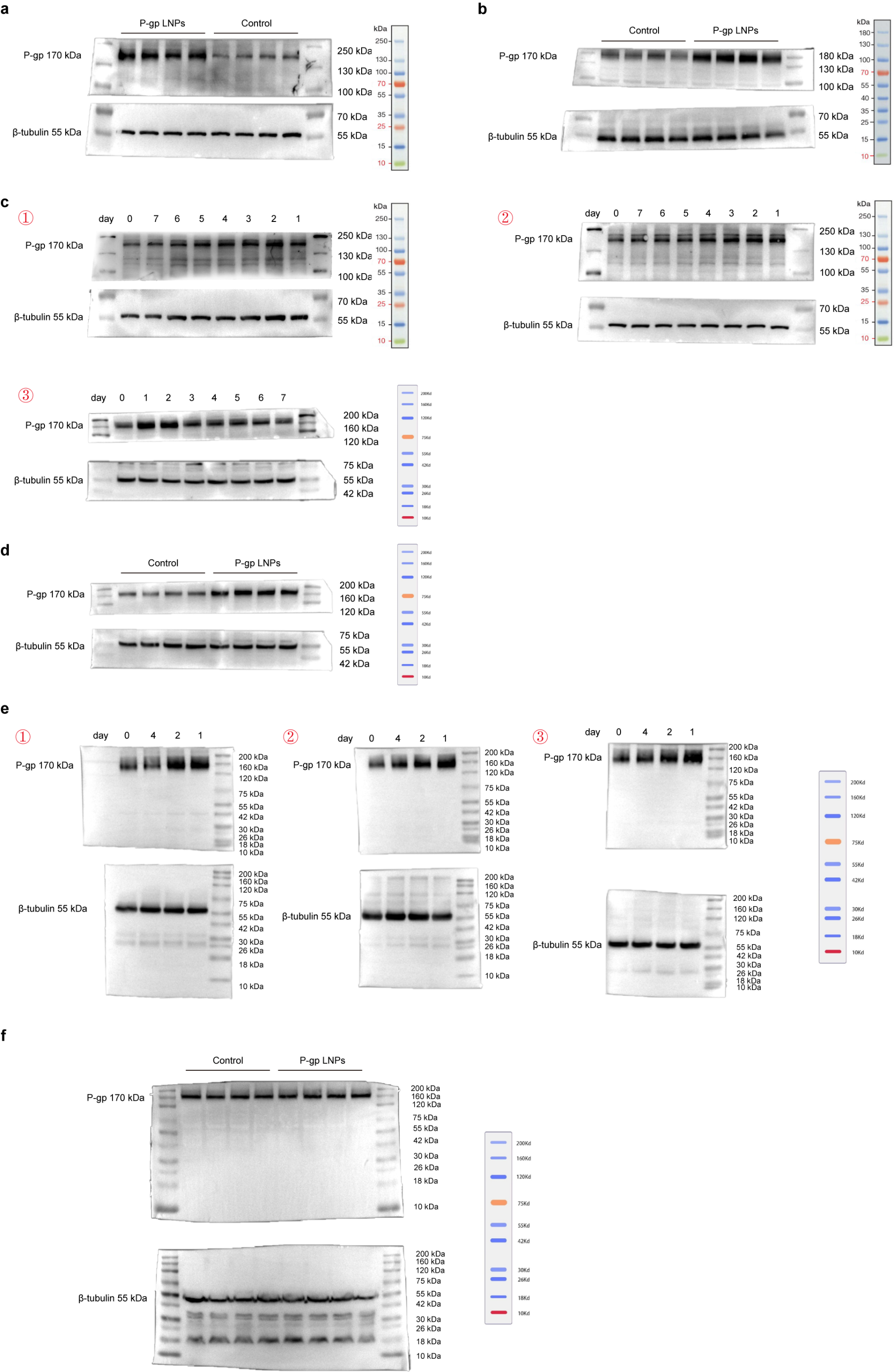

Supplementary figure S20**. Original blotting of western blot results. a,** Original blotting of western blot results for the analysis of P-gp expression in H9c2 cells with indicated treatments (Control and P-gp LNPs) (*n* = 4 independent biological samples). **b**, Original blotting of western blot results for the analysis of P-gp expression in mice heart tissues with indicated treatments (Control and P-gp LNPs) (*n* = 4 independent biological mice in each group). **c**, Original blotting of western blot results for the analysis of P-gp expression in H9c2 cells in 7 days after the treatment of P-gp LNPs (*n* = 3 independent biological samples). **d**, Original blotting of western blot results for the analysis of P-gp expression in PED cells with indicated treatments (Control and P-gp LNPs) (*n* = 4 independent biological samples). **e**, Original blotting of western blot results for the analysis of P-gp expression in mice heart tissues in 4 days after the intramyocardial administration of P-gp LNPs (*n* = 3 independent biological mice in each group). **f**, Original blotting of western blot results for the analysis of P-gp expression in mice tumor tissues after the intramyocardial administration of P-gp LNPs (*n* = 4 independent biological mice in each group).
